# Sphingolipid Control of Fibroblast Heterogeneity Revealed by Single-Cell Lipidomics

**DOI:** 10.1101/2021.02.23.432420

**Authors:** Laura Capolupo, Irina Khven, Luigi Mazzeo, Galina Glousker, Francesco Russo, Jonathan Paz Montoya, Sylvia Ho, Dhaka R. Bhandari, Andrew P. Bowman, Shane R. Ellis, Romain Guiet, Johannes Muthing, Bernhard Spengler, Ron M.A. Heeren, Gian Paolo Dotto, Gioele La Manno, Giovanni D’Angelo

**Author notes:** Correspondence to: G.D. ( / @GiO_DAngelo / epfl.ch/labs/dangelo-lab), G.L.M. ( / @GioeleLaManno / gioelelamanno.com).

## Abstract

Human cells produce thousands of lipids that impact a wide range of biological processes in ways we are only starting to characterize. The cellular composition in lipids changes during differentiation events and also varies across individual cells of the same type. Yet, the precise differences in lipid composition that directly affect cell phenotypes remain unknown. Here we have measured the lipidomes and transcriptomes of individual human dermal fibroblasts by coupling high-resolution mass spectrometry imaging to single-cell transcriptomics. We found that the cell-to-cell variation of specific lipid metabolic pathways contributes to the establishment of cell states involved in wound repair and in skin cancer growth. Sphingolipid composition defined fibroblast subpopulations while sphingolipid metabolic rewiring drove cell state transitions. These data uncover a role for cell-to-cell lipid heterogeneity in the determination of cell states and reveal a new regulatory component to the homeostasis and self-organization of multicellular systems.

## INTRODUCTION

The division of labor is a fundamental organizational principle of multicellular organisms. At a high scale, this is implemented through stable transcriptional programs that result in cell types: categories of cells with a specific role in the organism physiology. At a finer granularity, phenotypic heterogeneity across cells that belong to the same type results in cell states. These are metastable configurations within the landscape of phenotypes a cell type can explore [1]–[3]. Cell states have physiological significance, for example, they can provide a background priming on which diverging differentiation programs are triggered [4], or account for a dynamic repartition of labor in physiological processes [5].

With single-cell atlases made available for nearly all tissues, the existence and function of finer cell states are increasingly appreciated and the need for understanding the molecular determinants of state dynamics becomes more obvious [6]. While the role of transcriptional programs in defining cell states and guiding cell plasticity has been studied, much less is known about the function of cell metabolism [6]. Metabolic rewiring is inherent to cell fate transitions [7] and several metabolic switches that involve lipids occur in development [8]. Nonetheless, a few studies have investigated lipid composition at the single-cell level and the relevance of its variability. Thanks to these contributions, we know that several lipids are subjected to cell-to-cell variability as a result of microenvironmental cues [9], [10], cell cycle [11] or self-contained multi-stable circuits [12]. However, whether lipid metabolism has a role in the determination of cell states remains unclear.

Fibroblasts are an example of a plastic cell type that exists in multiple cell states [13], [14]. They synthesize and remodel the extracellular matrix and maintain the structural integrity of tissues [15]. Individual fibroblasts exhibit differences in gene expression that allowed the description of sub-populations, accounting for the fibrogenic, fibrolytic, and inflammatory activity of these cells [16]–[18]. Following tissue damage, fibroblasts experience phenotypic interconversion by adopting matrix secreting and inflammatory states [19], [20]. Changes in the proportions of these fibroblast subpopulations are associated with fibrosis and contribute to a tissue microenvironment permissive for cancer growth [21], [22]. Cell lineage, soluble factors, and microenvironment [13] have been implicated in the determination of fibroblast states [22], yet the molecular circuits that govern fibroblast heterogeneity and plasticity have not been fully clarified. For example, while specific lipids modulate the differentiation of stem cells in the skin [23], whether and how lipid metabolism participates in fibroblast state plasticity has not been addressed.

Recent technical developments have equipped mass spectrometry (MS) with enough sensitivity to enable single-cell lipidomics, thereby making the analysis of metabolic heterogeneity possible [24], [25]. In particular, matrix-assisted laser desorption/ionization mass spectrometry imaging (MALDI)-MSI provides extensive coverage of the lipid mass-to-charge-number (*m/z*) range, causes minimal fragmentation, and has reached a spatial resolution compatible with single-cell analysis, while maintaining good mass resolution and accuracy [26]–[33].

Here we coupled MALDI-MSI and single-cell mRNA sequencing (scRNA-seq) to investigate the metabolic and transcriptional heterogeneity of individual dermal human fibroblasts (dHFs). We found that multiple lipid compositional states exist in dHFs and that they correspond to transcriptional subpopulations. We also isolated the metabolic pathways that account for this correlation and found that specific lipids participate in cell-state determination. Changing lipid composition reprograms cells towards different cell states, and lipids modulate dHF response to instructive stimuli that drive their heterogeneity.

## RESULTS

### MALDI-MSI reveals the organizing principles of lipid heterogeneity

We devised a single-cell lipidomics approach by performing highly space-resolved (25-50 μm^2^pixel size) MAL-DI-MSI on low passage primary dHFs grown on glass coverslips and acquiring mass spectra in positive-ion mode (Fig. 1A). Lipid images (Fig. 1B) were extracted from raw data and lipid identity was attributed (see Methods) and validated by electrospray ionization liquid chromatography-mass spectrometry (ESI-LC/MS) and multiple reaction monitoring (MRM)-based lipidomics (Fig. 1A, S1A,B, and Table S1). Specific attributions were disambiguated by comparison with pure standards (Fig. S1C) and targeted LC-MS/MS (Fig. S1D). Based on the combination of these approaches, we associated an attribution score to lipid peaks (see Methods; Table S1). Overall, images of 205 annotated lipids were obtained, which account for a significant fraction of the dHF lipidome as detected by LC-MS.

**Figure 1.**
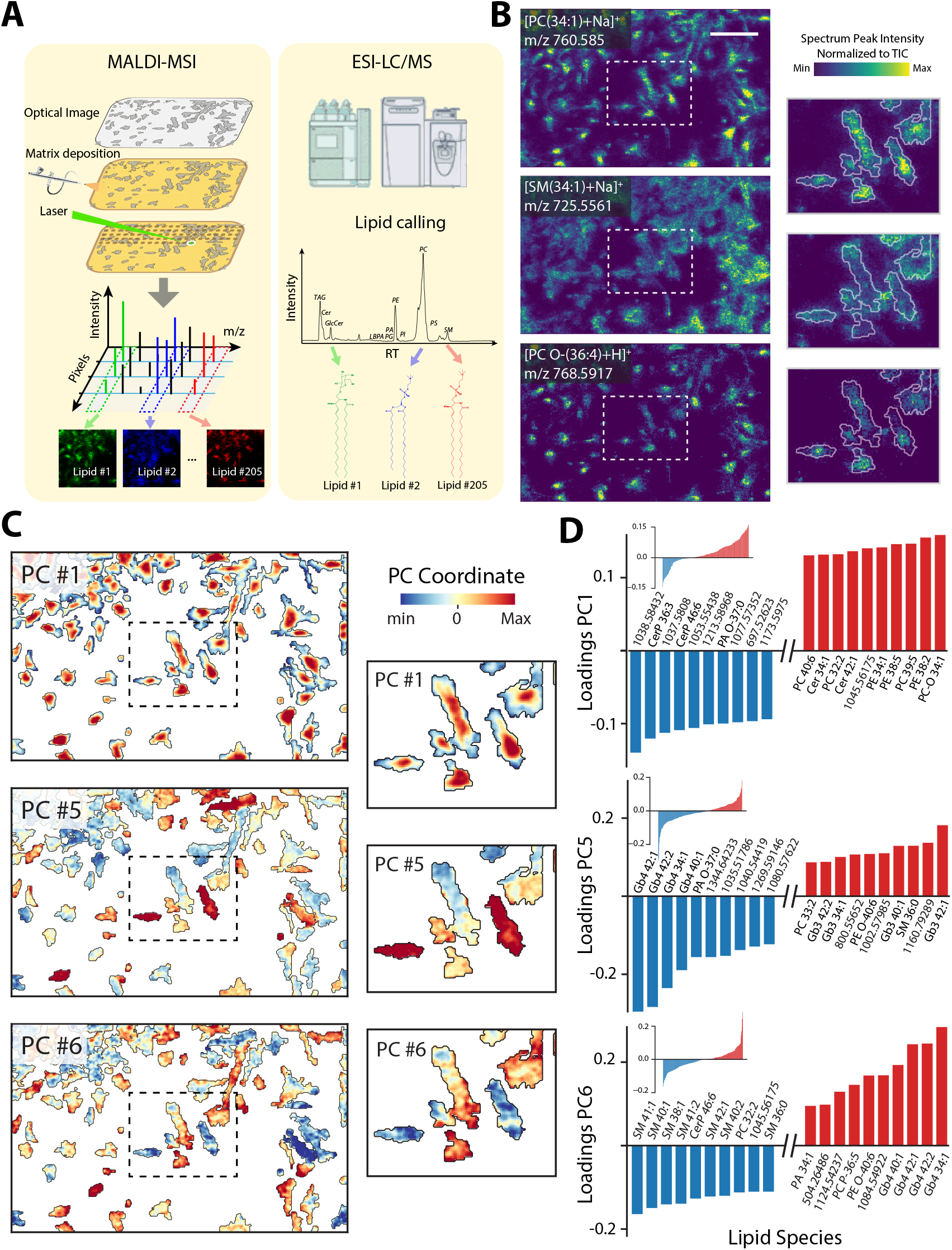
Single-pixel MALDI-MSI analysis on dHFs. (**A**) Schematic drawing of MALDI-MSI workflow. Cells were fixed, matrix was deposited and MAL-DI-MSI was performed by rasterizing the laser across a selected small area. For each spot, a mass spectrum was collected and mass images were obtained for each ion by plotting *m/z* intensity at the corresponding x,y coordinates (left panel). For peak identification, total lipid extracts were analyzed by ESI-LC/ MS (right panel). Lipids identified by ESI-LC/MS were then compared to the one obtained by MALDI-MSI (**B**) 50 μm^2^/ pixel ion images (354×218 pixels) of selected lipids recorded in positive-ion mode. Scale bar is 500 μm. (**C**) Images displaying at each location the PCA coordinate of each pixel. PC1, PC5 and PC6 values are displayed using a divergent colormap: positive coordinates in red and negative in blue. (**D**) Bar plots showing the contribution of the top ten lipids with higher (red) and lower (blue) loadings for each principal component. Miniatures show the entire distribution.

To learn how lipid composition varies spatially in dHF cultures, we considered the intensities of all the *m/z* peaks at each scanned location (i.e., pixel) and performed a multivariate analysis. Principal component analysis (PCA) revealed that 95% of the pixel-to-pixel variability is explained by eight principal components (PCs) (Fig. S1E). The *in situ* visualization of the PC coordinates corresponding to each pixel delineated distinct distribution patterns for different groups of lipids (Fig. 1C).

PC1 coordinates changed moving from the inner part of the cell towards the cell periphery. Inspecting the contributions to PC1, we found high loadings corresponding to both glycerolipids and sphingolipids (Fig. 1D), confirming that this axis captures fundamental differences in lipid composition of the perinuclear and peripheral cell membranes. In contrast to what was observed for PC1, where the trend was reproduced similarly across cells, PC2-8 coordinates distributed differently among cells, with some cells displaying exclusively positive or negative pixels (Fig. 1C). When we asked which features accounted for these axes of cell-to-cell variation we found lipids belonging to the sphingolipid pathway (i.e., ceramides [Cer]; sphingomyelins [SM]; hexosylceramides [HexCer]; trihexosylceramides [Gb3]; and globosides [Gb4]) (Fig. 1D and Fig. S1E). This confirms previous observations from our group and others, concerning the cell-to-cell variability of selected sphingolipids [9], [12] and extends them to most of the lipid species in the sphingolipid pathway.

Overall, this analysis revealed two coexisting axes of lipid variation in dHFs. One axis mirrored intracellular organization [34] while the other captured lipid-related inter-cellular heterogeneity [35].

### Single-cell analysis reveals lipid coregulation

The multivariate analysis we performed suggested that specific classes of lipids are involved in the cell-to-cell variability observed. To evaluate the biological significance of our finding, we asked whether this heterogeneity could be attributed to the general activation of a metabolic process or to the regulation of specific enzymatic reactions. We used optical images to guide cell segmentation and transferred it onto the MS images to obtain a total of 257 single-cell lipidomes from three independent MAL-DI-MSI recordings (Fig. 2A). After data normalization and batch correction, the cell-to-cell variability associated with individual lipid species was summarized by computing their coefficient of variation across the cell population (CV; the ratio of standard deviation over mean [σ/μ] of the normalized intensity values). The obtained values were used to rank lipids according to their decreasing CV. Sphingolipids populated the top CV ranking positions indicating again that the sphingolipid pathway (Fig. S2A) is subjected to major cell-to-cell variability (Fig. 2B). Nevertheless, it remained to be determined whether the entire sphingolipid metabolic pathway is coordinately modulated or different sphingolipid subsets are controlled independently in different cells.

**Figure 2.**
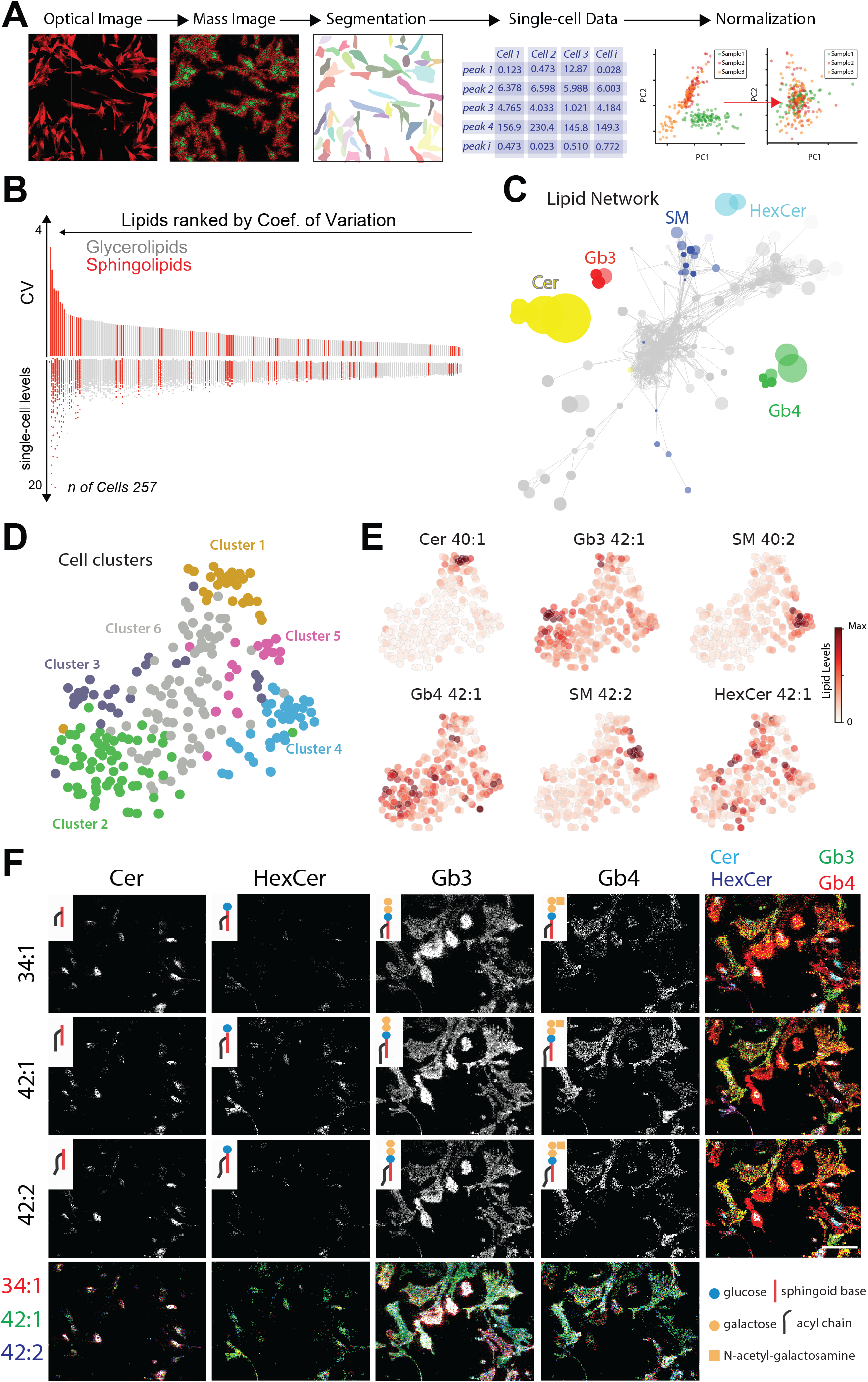
Single-cell lipidomics analysis. (**A**) Schematic of the approach used for single-cell analysis of MALDI-MSI data. Confocal micrographs were used as guides to segment cells out of mass images and single-cell ion abundance computed as the total total ion current (TIC)-normalized peak intensity. Different acquisitions were combined after ComBat batch correction [59]. (**B**) Barplot showing the coefficient of variation (CV) of lipids computed across 257 cells. Lipids were ranked by CV and color coded according to their lipid class (sphingolipids in red, glycerolipids in grey). Single-cell lipid levels are shown in the bottom part of the plot. (**C**) A lipid co-variation network. Nodes represent individual lipids, size is proportional to the CV, color is according to lipid class. Edges connect two lipids where the correlation coefficient is > 0.85. (**D**) t-SNE embedding of the single-cell lipidomics data. Cells are colored by the clusters defined by hierarchical clustering. (**E**) t-SNE embedding colored by the abundances of sphingolipids. (**F**) Mass images showing the spatial distribution of sphingolipid precursors (Cer, HexCer) and complex sphingolipids (Gb3 and Gb4). Miniatures in the top left corner of each image depict a simplified schematic of the lipid structure (cf. Figure S2A). Scale bar is 500 μm.

With this aim in mind, we assessed which lipid species co-vary across cells. A pairwise lipid-lipid correlation (Pearson’s R) matrix was computed and summarized as a network where lipid species were represented as nodes and connected when lipids strongly co-varied (R > 0.85) (Fig. 2C). Interestingly, while phospholipid species did not form biochemically meaningful cliques, sphingolipids were clustered in groups consisting of compounds bearing the same headgroup (i.e., OH with ceramides; Hexose with HexCers; Trihexose with Gb3 species; and N-acetyl-hexose-trihexose with Gb4 species) but with different ceramide backbones (mostly 34:1, 40:1, 42:1, and 42:2) (Fig. S2A). This indicates that ceramide processing is more cell-to-cell variable than the production of different ceramides by dedicated ceramide synthases. Accordingly, the relative abundance of lipids sharing the same headgroup were found to be more correlated than those sharing the same backbone (Fig. S2B).

We next asked whether co-varying sphingolipids define cell sub-populations. Single-cell lipidomes were used to group cells according to their lipid composition (see methods for details) resulting in distinct cell clusters (Fig. 2D, Fig. S2C). When the levels of sphingolipids were considered, we observed that certain species (i.e., Ceramides, HexCers, Gb3s and Gb4s) were enriched in specific cell clusters, suggesting that dHFs exist in distinct sphingolipid metabolic states (Fig. 2E-F).

### Sphingolipids define dHF lipotypes *in vitro* and *in vivo*

To confirm lipid variability in dHFs we stained cells with fluorescently-labelled bacterial toxins that recognize different sphingolipid headgroups: Shiga toxin 1a (ShTxB1a) binds to Gb3 [36], Shiga toxin 2e (ShTxB2e) binds to Gb3 and Gb4 [37], and Cholera toxin (ChTxB) binds the ganglioside GM1 [38]. We found that toxins stained dHFs in a cell-specific fashion, with a pattern reminiscent of the variability observed by MALDI-MSI (Fig. 3A, B). Treatment with Fumonisin B1 [FB1] [39], and D-threo-1-phenyl-2-decanoylamino-3-morpholino-1-propanol [D-PD-MP][40]), two inhibitors of sphingolipid production, blocked toxin binding indicating that toxins are a faithful proxy for dHFs sphingolipid composition in our setting (Fig. S3A).

**Figure 3.**
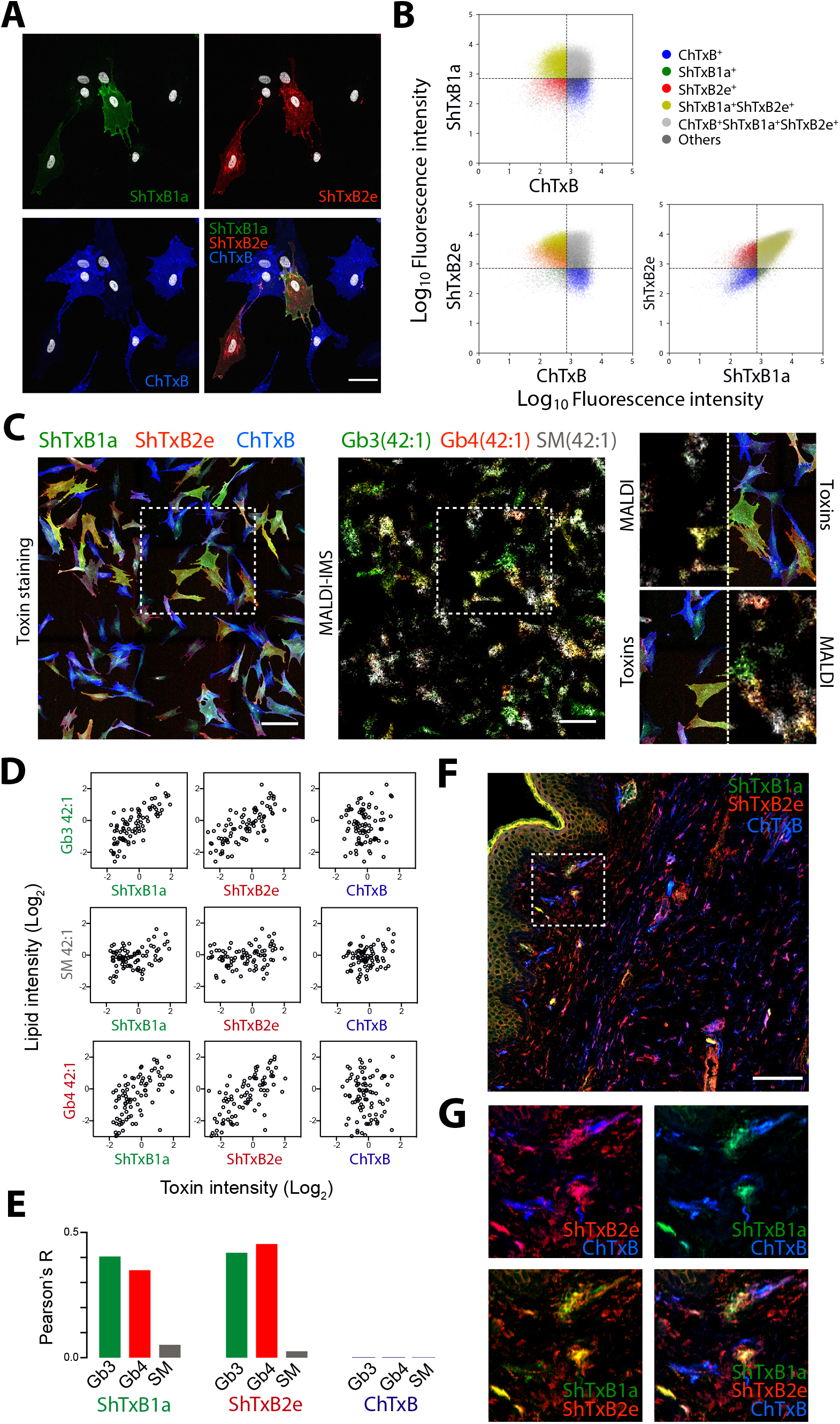
Identification of dHF lipotypes by MALDI-MSI and toxin staining. (**A**) Confocal micrographs showing cells stained using bacterial toxins ShTxB1a (green), ShTxB2e (red), ChTxB (blue), and Hoechst (grey) for nuclei. Scale bar is 50 μm. (**B**) Cytofluorometric analysis of dHFs with bacterial toxins. Scatter plots of fluorescence intensity values are shown for each toxin. Populations are colored according to their lipid composition. Unstained cells were used as negative control to determine the gates; n = 50’799. (**C**) Side-by-side comparison of toxin staining and MALDI-MSI acquisition on the same cells. First, cells were stained with bacterial toxins ShTxB1a (green), ShTxB2e (red) and ChTxB (blue) and images acquired by confocal microscopy (left panel). Then, MALDI-MSI measurements (25 μm^2^/pixel) were performed on the same cells (center panel). Mass images (320×320 pixels) of complex sphingolipids [SM (42:1), Gb3 (42:1) and Gb4 (42:1)] showed good correlation with the confocal image as shown in the inset. Scale bar is 200 μm. (**D**) Scatterplots comparing toxin staining intensity and lipid abundances as determined by MALDI-MSI. Data were log2 transformed and centered. (**E**) Barplot summarizing the Pearson’s coefficient (R) for each toxin-lipid couple shown in (D); n= 88. (**F**) Confocal micrographs of human foreskin tissue section stained with bacterial toxins ShTxB1a (green), ShTxB2e (red) and ChTxB (blue) and analysed by confocal microscopy. Scale bar is 100 μm. (**G**) Insets of skin tissue section show cell-to-cell sphingolipid variability in the dermis.

As a further validation, we directly compared toxin staining and MALDI-MSI. To this aim, dHFs were first fixed and stained with ShTxB1a, ShTxB2e and ChTxB and then imaged by MALDI-MSI. A close correspondence was found between the two imaging methods (Fig. 3C). ShTx-B1a staining correlated best with Gb3 levels and ShTxB2e staining correlated well with Gb3 and Gb4 levels, whereas neither correlated with SM levels. ChTxB staining, our proxy for the levels of GM1 [38], a sphingolipid not detected by MALDI-MSI in positive-ion mode, did not correlate with any of the sphingolipids detected by mass imaging (Fig. 3D,E).

We then asked whether the discovered cell-to-cell variability is specific to the cell line and the *in vitro* conditions or it is a generalized phenomenon. To this aim, we analyzed four dermal fibroblast lines derived from unrelated healthy individuals by toxin staining and cytofluorometry. All the tested fibroblast lines displayed an analogous pattern of distribution of the sphingolipids across cells (Fig. S3B). Importantly, we toxin-stained dermal cells in the context of skin biopsies and found that also in their physiological context dermal fibroblasts display remarkable cell-to-cell sphingolipid heterogeneity (Fig. 3F,G and Fig. S3C,D).

These data confirm more broadly and *in vivo* that dHFs exist in different metabolic states characterized by their sphingolipid composition. Hereafter we refer to these lipid metabolic states as lipotypes.

### Lipotypes mark specific cell states

To test if specific cell states are associated with the different lipotypes, we performed single-cell RNA sequencing (scRNA-seq) for a total of 5652 dHFs. Uniform manifold approximation and projection (UMAP) embedding was computed on the gene expression profiles and 17 cell clusters were identified by the Louvain algorithm (Fig. 4A). When the relationship among these 17 groups was investigated we found they could be grouped into five categories, each related to a different biological process: proliferation, pro-inflammatory cytokine secretion (inflammatory), pro-fibrotic secretion (fibrogenic), extracellular matrix remodeling (fibrolytic) and pro-angiogenic factor secretion (vascular) (Fig. 4B, S4A). A further group represented bona fide basal-state fibroblasts (basal).

**Figure 4.**
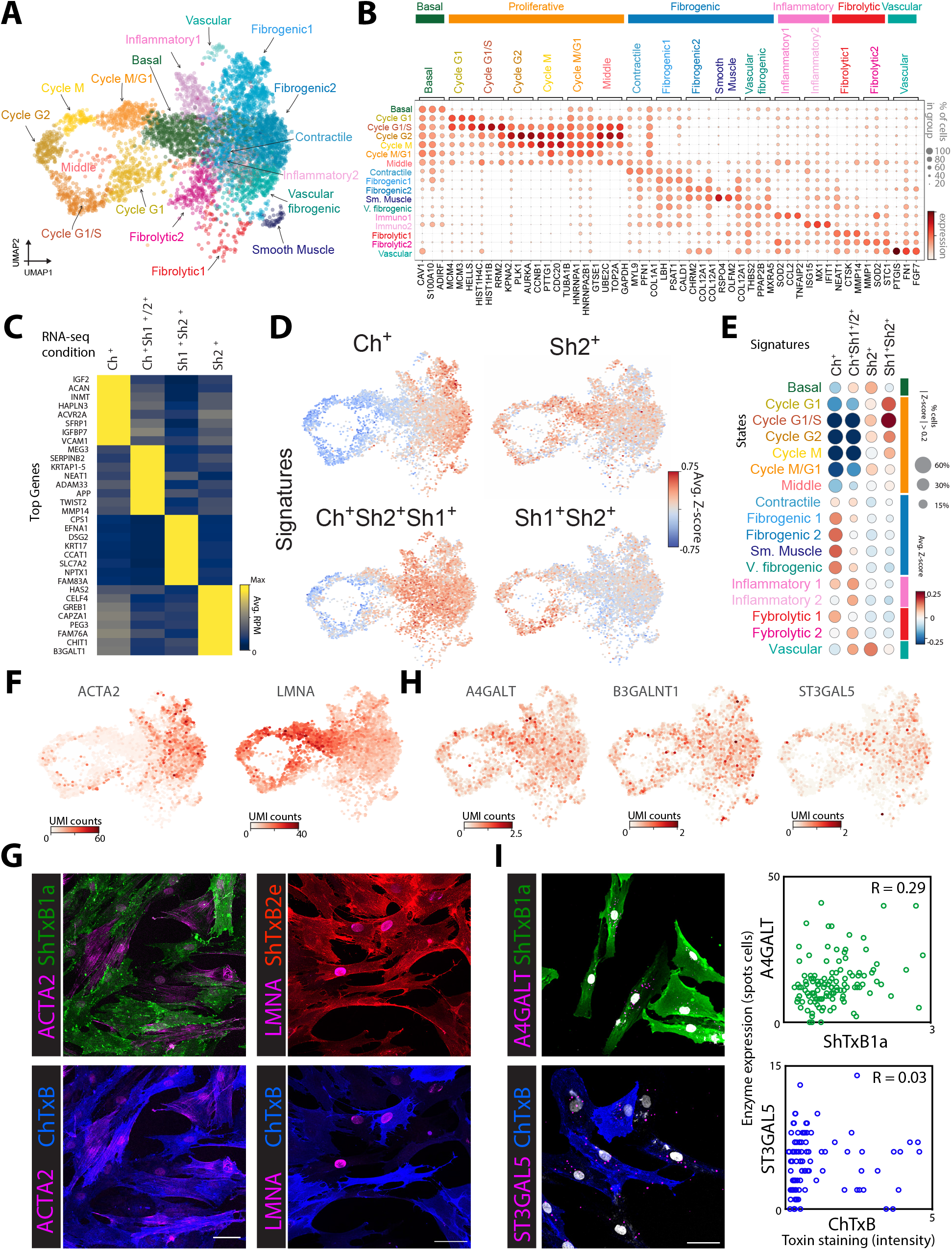
Lipotype mapping to transcriptional cell states. (**A**) UMAP embedding analysis of scRNA-seq of 5652 individual dHFs colored by the assigned cluster. (**B**) Gene expression dot plot of these genes is shown. Marker genes for each cluster were identified using the Wilcoxon test. (**C**) Heatmap reporting the average gene expression of enriched genes for each of the FACS-sorted lipotype populations (bulk RNA-seq data). For each lipotype the top eight genes, ranked by fold change, are shown. (**D**) UMAP embedding colored by the different lipotype gene signature scores. 250 top differentially expressed genes were used to calculate the signature score. (**E**) Dot plot colored by the average lipotype Z-score of cells of the different clusters. Size of the dots represent the number of cells with magnitude of the score >0.35. (**F**) UMAP embedding colored by the expression of the two canonical markers for fibrogenic, *ACTA2*, and basal cell states, *LMNA*. (**G**) Confocal images of cells stained with ShTxB1a (green), ShTxB2e (red) and ChTxB (green) and counterstained by antibodies against ACTA2 and LMNA (magenta). Scale bar is 100 μm. (**H**) UMAP embedding is colored by the expression of *A4GALT* (Gb3S), *B3GALN*T1 (Gb4S) and *ST3GAL5* (GM3S). (**I**) Left, representative confocal microscopy images of correlative mRNA-FISH/fluorescence toxin staining using *A4GALT* and *ST3GAL5* (magenta) probes and ShTxB1a (green) and ChTxB (blue). Nuclei were labelled with Hoechst (grey). Scale bar is 50 μm. Right, scatterplot showing level of expression against toxin fluorescence. Pearson’s correlation coefficient is indicated. Quantification on 120 and 96 individual cells for *A4GALT* and *ST3GAL5* respectively.

Next, to link the expression-defined subtypes with those defined by sphingolipids, we isolated dHFs according to their lipotypes by fluorescence activated cell sorting (FACS) and performed RNA sequencing on the different sorted samples. We isolated ChTxB^+^/ShTxB1a^-^/2e^-^(henceforth ChTxB^+^), ShTxB2e^+^/ChTxB^-^/ShTxB1a^-^(ShTx-B2e^+^), ShTxB1a^+^/2e^+^/ChTxB^-^(ShTxB1a/2e^+^), and ShTx-B1a^+^/2e+/ChTxB^+^ (triple positive) cells (Fig. 4C, S4B). Genes upregulated in the different lipotypes were extracted and used to compute gene signatures scores on the single-cell dataset. The four lipotype signatures mapped to distinct UMAP areas that corresponded to the major transcriptional clusters (Fig. 4D, E). Triple-positive cells corresponded to inflammatory, fibrolytic, and vascular fibroblasts, ShTxB1a/2e^+^ to proliferating cells, ShTxB2e^+^ to basal state fibroblasts, and ChTxB^+^ to ‘fibrogenic’ fibroblasts, suggesting that specific membrane lipid compositions are associated with prevalent cell states. To verify this finding, we co-stained dHFs with toxins and markers for the different clusters. The stainings revealed a good correlation between ChTxB and smooth muscle actin (encoded by the gene *ACTA2*; fibrogenic marker) and ShTxB2e and laminin A (encoded by the gene LMNA; basal and proliferative marker) altogether indicating that lipotypes are markers of dHF cell states (Fig. 4F, G).

We then asked whether lipotypes are the result of cell state-specific transcriptional programs that involve lipid metabolising enzymes. Surprisingly, when the expression of genes encoding sphingolipid enzymes and accessory factors was visualised on the UMAP embedding none of them showed a cell-state specific localisation (Fig. 4H and S4C). To test this more directly we set up a correlative analysis combining toxin staining and mRNA fluorescence in situ hybridization (FISH). We probed in the same cell the expression of *ST3GAL5* encoding GM3 Synthase [GM3S] and *A4GALT* encoding Gb3 Synthase [Gb3S]) (Fig. S4D) and their lipid products through toxins ChTxB and ShTxB1a (Fig. 4I). When toxin staining intenisity was considered along with FISH counts, we observed either no or weak (R=0.29) correlation of the two readouts, suggesting that single-cell sphingolipid composition is largely determined by post-transcriptional mechanisms.

Thus, lipotypes are associated with cell states, yet cell states are not endowed with transcriptional programs that would account for the lipotypes they are associated with. This raises the question of whether lipotypes are causally upstream of cell states, or in other words, whether lipid composition influences cell-to-cell transcriptional heterogeneity.

### Sphingolipid composition influences cell states

We treated dHFs with the ceramide synthase inhibitor FB1 that blocks the production of sphingolipids (Fig. S2C) and performed scRNA-seq. When FB1-treated dHFs were integrated in the same transcriptional embedding along with control cells, they displayed a different distribution across states (Fig. 5A and S5A). FB1-treated cells were more frequently associated with fibrolytic (from 6% in CTRL to 23% in FB1 treated) and vascular (from 0.6% to 1.3%), and less with fibrogenic (from 48% to 40%) and inflammatory (from 9% to 6%) (Fig. 5A and S5B) states.

**Figure 5.**
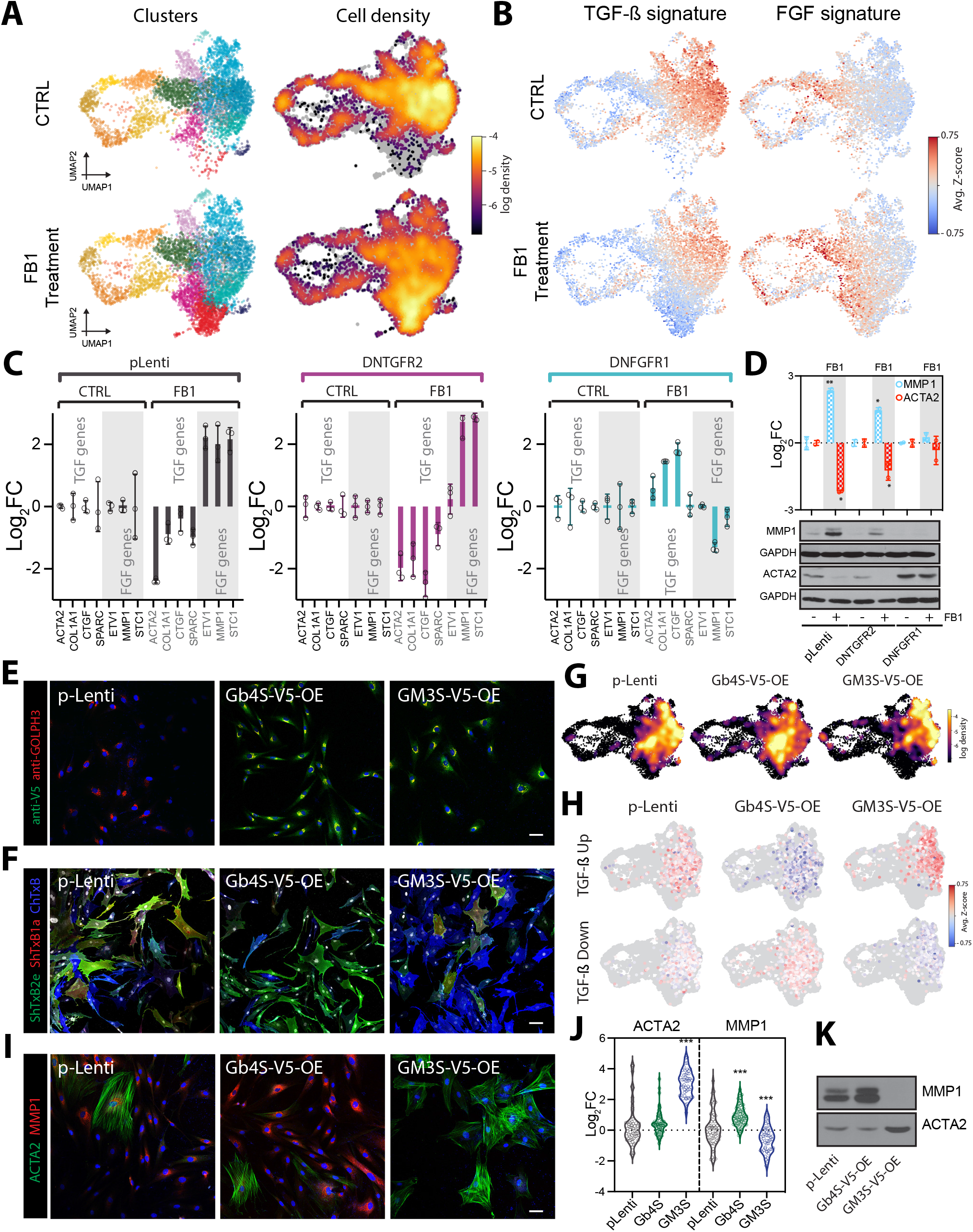
Effect on cell states of sphingolipid perturbations. (**A**) Left, UMAP embedding of the scRNAseq data for the CTRL (5652 individual dHFs) and FB1 treated cells (6546 individual dHFs). Cells colored by their assigned cluster are shown in the left panel. Right, density maps of CTRL and FB1 treated cells mapped in the UMAP space. (**B**) TGF-β and FGF2 pathway activation signatures in the CTRL (upper panel) and FB1 treated sample (lower panel) overlayed onto the UMAP embedding. (**C**) Barplots of a qPCR quantifying the mRNA levels of TGF-β and FGF2 target genes in CTRL, DNTGFR2 and DNFGFR1 cells treated with 25 µM FB1. Data are shown as log2 fold change over untreated cells (n=3; data are means ± StDev). (**D**) Western blot and their quantification, for cells treated as in (**C**). Data were normalized against GAPDH (n=2; data are means ± StDev; *p <0.05, **p <0.01 [Student’s t-test]). (**E**) Confocal images of the overexpressing cell lines stained with antibodies against V5 protein tag (green) and GOLPH3 (red). Scale bar is 50 μm. (**F**) Confocal images of the overexpressing cell lines stained with bacterial toxins ShTxB1a (red), ShTxB2e (green) and ChTxB (blue). Scale bar is 50 μm. (**G**) Cell density plot of single-cell expression profile of the OE-dHFs cells mapped by similarity onto the UMAP projection in (A). (**H**) TGF-β and FGF2 pathway activation response gene signatures overlayed to the OE-dHFs cells mapped to the UMAP as in (**G**). (**I**) Confocal images of cells stained with antibodies against ACTA2 (green) and MMP1 (red). Scale bar is 100 μm. (**J**) Quantification of the immunofluorescence measurements. Data were scaled to the median and shown as log2 fold change over control in individual cell (CTRL n= 58; GM3S-OE n= 49; Gb4S-OE n= 81;***p <0.001 [Student’s t-test]). (**K**) Western blot of the OE-dHFs cells with antibodies recognizing ACTA2 and MMP1.

Upon examination of the transcriptional changes triggered by FB1, we noticed that they involved genes, such as *MMP-1, MMP-2, MMP-14*, and *STC1*, that are either induced by Fibroblast Growth Factor 2 (FGF2) or repressed by Transforming Growth Factor Beta (TGF-β) signalling [41] (Fig. S5C). FGF2 and TGF-β drive opposing transcriptional programs involved in the determination of alternative fibroblast states [41]. Intrigued by the possible involvement of those signaling pathways in the effect of FB1, we systematically analyzed the single-cell signatures for the enrichment of one or the other pathway. We computed transcriptional signature scores corresponding to the activation of FGF2 or TGF-β pathway in dHFs (See Methods). Genes upregulated by FGF2 were preferentially expressed in the FB1-treated cells and in the fibrolytic population, while genes upregulated by TGF-β were more expressed in control cells and in the fibrogenic population (Fig. 5B), suggesting that FB1 (and thus sphingolipid deprivation) either promotes FGF2 or suppresses TGF-β signaling. qPCR and immunofluorescence analysis on selected markers confirmed this and extended the observation to the treatment with D-PDMP (Fig. S5D,E).

To untangle which of these two axes is affected by sphingolipid depletion we challenged control and FB1-treated dHFs with increasing amounts of FGF2 or TGF-β1. Sphingolipid depletion did not inhibit fibroblast response to TGF-β while it sensitized cells to FGF2 (Fig. S5F). As a more direct test, we used dHF lines expressing dominant negative versions of either FGF receptor 1 (DNFGFR1) or TGF-β receptor 2 (DNTGFBR2) [41] where the FGF2 and TGF-β signaling are blocked, respectively. We treated these cells with FB1 and evaluated their response by real-time qPCR and Western blotting. The results revealed that, while FB1 clearly upregulated FGF2 target genes and downregulated TGF-β target genes in both control (pLenti) and DNTGFBR2 fibroblasts, it was ineffective in inducing the same transcriptional changes in DNFGFR1-dHFs (Fig. 5C,D). These data show that FB1 requires an active FGF2 pathway to exert its transcriptional effect and, thus, that sphingolipids influence FGF2 transcriptional programs possibly modulating FGFR activity at the plasma membrane.

### Different sphingolipids drive specific cell-state switches

FB1 treatment deprives cells of most sphingolipids [39],; hence, this treatment does not inform on how the individual lipid species associated with the cell states influence FGF2 signaling. Thus, we generated dHFs overexpressing either GM3S or Gb4S, the two enzymes driving alternative sphingolipid processing pathways (Fig. 5E). These over-expressing (OE) cells displayed the expected changes in sphingolipid composition with GM3S-OE dHFs being largely ChTxB^+^ and ShtxB1a/2e^-^, and Gb4S-OE dHFs being prevalently ShtxB1a/2e^+^ and ChTxB^-^(Fig. 5F and S6A).

GM3S-OE and Gb4S-OE lines were then analyzed by scR-NA-seq to test the impact of lipotype change on cell state. Interestingly, GM3S-OE and Gb4S-OE dHFs populated two distinct transcriptional regions (Fig. 5G). Gb4S-OE dHFs were more associated with inflammatory and fibrolytic states, while GM3S-OE dHFs were for the major part in a fibrogenic state (88%) and almost never in inflammatory or fibrolytic states (Fig. S6B). Gene expression analysis confirmed this transition: *COL12A1* and *VCAN* markers of fibrogenic state were significantly upregulated in GM3S-OE and downregulated in Gb4S-OE cells, while fibrolytic and inflammatory markers *MMP1* and *CCL2* were upregulated in Gb4S-OE cells and downregulated in GM3S-OE cells (Fig. S6C).

When FGF2 and TGF-β signatures were mapped onto Gb4S-OE and GM3S-OE dHFs, we observed that GM3S OE fostered the TGF-β transcriptional program, while for Gb4S-OE we revealed the opposite trend (Fig. 5H). The effect on the FGF2 program was more difficult to appreciate as the expression signature dominates in actively proliferating cells (Fig. S6E). Nonetheless, immunofluorescence and Western blot experiments showed that while GM3S-OE dHFs were almost uniformly ACTA2^+^/ MMP1^-^, Gb4S-OE dHFs display high MMP1 levels (Fig. 5I-K). Altogether, this evidence suggests that changes in sphingolipid composition drive cell state switches by specifically influencing cell response to instructive stimuli.

### Sphingolipids integrate into regulatory circuits involved in cell-state determination

Our results established that sphingolipids participate in the determination of the cell states with which they are associated and that they interact with gene expression programs downstream of FGF2 and TGF-β pathways. To complete our understanding of the directionality of those interactions we sought to determine whether FGF2 and TGF-β pathway modulation can also affect sphingolipid metabolism. In order to test this possibility, we performed toxin staining and cytofluorimetric analysis in control, DNFGFR1, and DNTGFBR2 dHFs. The inactivation of the TGF-β pathway had little effect on the toxin binding pattern (i.e., a slight decrease in ChTxB^+^ cell populations), while interruption of the FGF2 signaling caused a transition of the dHFs to a ChTxB^+^ state (Fig. 6A-C). Accordingly, when the sphingolipid synthesis was analyzed by [H^3^]-sphingosine pulse experiments, the production of the Gb3 was significantly reduced in DNFGFR1 dHFs (Fig. 6D).

**Figure 6.**
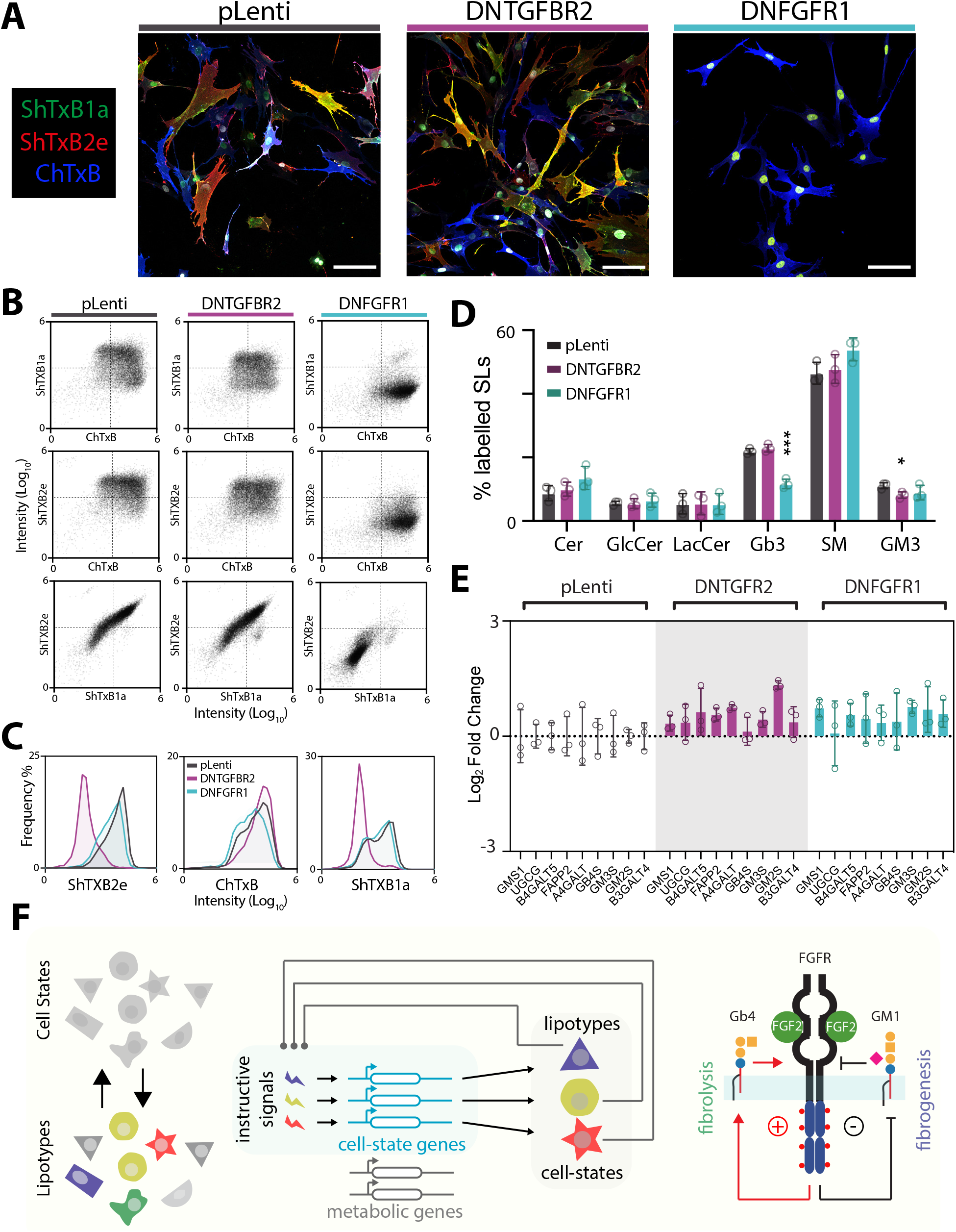
Effect of perturbations of signaing on lipotypes. (**A**) Confocal images of pLenti, DNTGFBR2 and DNFGFR1. Cells were stained with bacterial toxins ShTxB1a (green), ShTxB2e (red), ChTxB (blue) and Hoechst for nuclei. Scale bar is 100 μm. (**B**) Cytofluorimetric scatters of pLenti, DNTGFR2 and DN-FGFR1 cells stained with bacterial toxins. Unstained cells were used as negative control and to determine the gates (pLenti n=12’214; DNFGFR1 n=12’855; DNTGFR2 n=12’357). (**C**) Histograms of the frequency distribution of the binding of different toxins for each condition. (**D**) Barplots of dHFs sphingolipid synthesis activity as determined by a 2 hours pulse labelling with [H^3^]-D-Sphingosine. The percentage of total radioactivity associated with SM, Cer, GlcCer, LacCer, Gb3, and GM3 in the different cell lines was quantified after lipid extraction and HPTLC separation. (n=3; data are means ± StDev; *p <0.05, ***p <0.001 [Student’s t-test]). (**E**) Barplots of a qPCR quantifying genes encoding SLs synthetic enzymes for pLenti, DNTGFR2 and DNFGFR1 cells. mRNA levels were evaluated in the three cell lines. qPCR data are shown as log2 fold changes over control (n=3; data are means ± StDev). (**F**) Schematic representation of the model for the role of lipotypes on cell-state determination consistent with our data. Left panel, lipotypes correspond to dHF cell states. Middle panel, cell states/lipotypes are determined by signaling pathways that are influenced by the lipid composition of individual cells. Right panel, FGF2 binds FGFR leading to the prevalent production of Gb3/Gb4 over GM1. To close the circuit GM1 negatively regulates FGFR, while Gb3 and Gb4 activate FGFR in a positive feedback loop. This circuit generates a bistable system where cells can be either Gb3/Gb4^+^ leading to a more fibrolytic state or GM1^+^ leading to a more fibrogenic state.

Interestingly, when looking at the expression of the genes that encode sphingolipid synthetic enzymes in DNF-GFR1 and DNTGFBR2 we did not find any significant modulation (Fig. 6E). Thus, we concluded that the FGF2 signaling regulates the sphingolipid synthetic system to promote the production of globo-series sphingolipids and to repress the expression of gangliosides by modulating the pathway at a post-transcriptional level.

These data show that multiple populations of dHF coexist in culture, and bear proliferative, inflammatory, fibrogenic, and fibrolytic properties. These populations are characterized by distinct lipotypes and are influenced by opposing signaling pathways. Sphingolipids (i.e., GM1 and Gb4) modulate these signaling pathways by inhibiting or fostering the FGF2 signaling. In turn, FGF2 signaling counteracts GM1 production by sustaining the alternative metabolic pathway leading to the production of Gb3 and Gb4. This regulatory wiring leads to a circuit where its operation accounts for the phenotypic heterogeneity of dHFs (Fig. 6F).

## DISCUSSION

Lipids participate in energy metabolism, are fundamental for the assembly of biological membranes, function as signaling molecules, and interact with proteins to influence their activity and intracellular distribution [42]. Certain lipids are then involved in biological processes that are either cell-type or developmental stage specific [43]–[47]. Still, whether specific lipid metabolic configurations influence cell identity, or are merely a consequence of differentiation programs was poorly understood.

In this study, we considered the case of dHF heterogeneity as a prototypical example of dynamic cell identity [14], [21]. dHF subtypes represent cell states that can plastically interconvert as a result of environmental cues and instructive signals [13], [48]–[50]. By coupling MAL-DI-MSI analysis with scRNA-seq we uncovered a two-way causal relationship between dHF cell states and their lipid composition.

Cell states are intermediates in the process of cell differentiation where state switches precede fate commitments [4]. Lipidome remodeling can thus be an early driver in the establishment of cell identity and possibly promoting symmetry breaking events [51]. Nonetheless, whether *in vivo* lipid metabolic fluctuations can prime cells towards discrete transcriptional states remains to be determined. Irrespective of this, our data suggest that discrete lipid compositions cause cells to respond differently to identical stimuli. While this is likely the consequence of lipid modulation of plasma membrane receptors [52]–[54] the mechanistic details and the specificity rules of such modulation are unknown. Addressing these points will provide a new perspective to our understanding of the role of cell metabolism in multicellularity.

This study also demonstrates the potential of single-cell lipidomics to reveal biological phenomena that cannot be studied using classical lipidomics. This suggests that single-cell MS-based lipid analysis coupled to other single-cell techniques will allow us to explore still uncharted domains of biology. In our case, this approach permitted us to characterize a role for sphingolipids in dHF heterogeneity. This heterogeneity plays a major role in skin homeostasis and wound repair. It is also observed in cancer-associated fibroblasts where different proportions of inflammatory, immune or fibrogenic fibroblasts influence tumor growth and metastasization [55]–[58]. Our data indicate that cell-to-cell heterogeneous sphingolipids (Gb3, Gb4, and GM1) integrate into the circuits that drive the activation of dHFs through FGF2 signaling and thus suggest that pharmacological modulation of sphingolipid metabolism holds promise for the prevention of fibrotic lesions and to target the tumor microenvironment.

## ACKNOWLEDGEMENTS

We thank Bart Deplancke, Parashuraman Seetharaman, Domenico Russo, Charna Dibner, Yussuf Hannun, Chiara Luberto and Sten Linnarsson for critically reading the manuscript. We thank Krisztian Homicsko for scientific advice and fibroblast reagents. G.L.M and I.K. were supported by the Swiss National Science Foundation grants CRSK-3_190495 and PZ00P3_193445. G.DA. acknowledges financial support from the Swiss Cancer League (KFS-4999-02-2020), EPFL institutional fund, Kristian Gerhard Jebsen Foundation and from the Swiss National Science Foundation (SNSF) (grant number, 310030_184926).

## AUTHOR CONTRIBUTIONS

L.C. developed the idea, conducted the experiments and wrote the manuscript; I.K. participated in the development of the idea and conducted image and scRNA-seq analyses and experiments; L.M. supported L.C. in experiments on DN fibroblast lines; G.G. generated GM3S and Gb4S OE fibroblast lines, F.R. performed initial lipid data analysis, J.P.M. performed targeted and untargeted lipidomics, S.H. provided technical assistance, A.P.B., S.R.E. for assistance with MALDI-MSI, R.G. for FISH analysis, J.M. provided ShTxB2e toxin, D.R.B. and B.S. contributed on AP-SMALDI10 and AP-SMALDI5 AF experiments, R.M.A.H. provided assistance with MALDI-MSI data treatment, G.P.D. contributed with discussion and reagents for dHF heterogeneity, G.L.M. and G.DA. developed the idea, designed and supervised the entire project, analyzed the data, and wrote the manuscript.

## DATA AVAILABILITY

Lipid images are available on the metaspace website at https://metaspace2020.eu/api_auth/review?pr-j=09e813fe-6b8f-11eb-96db-1b87e6b0215b&token=-j43VwRscgJF-.

Sequencing data are available on GEO database, under acession GSE167209.

Analysis notebooks are available at https://github.com/lamanno-epfl/capolupo2021.

mRNA-FISH analysis script is available at https://doi.org/10.5281/zenodo.4548877.

## SUPPLEMENTAL FIGURES

**Figure S1.**
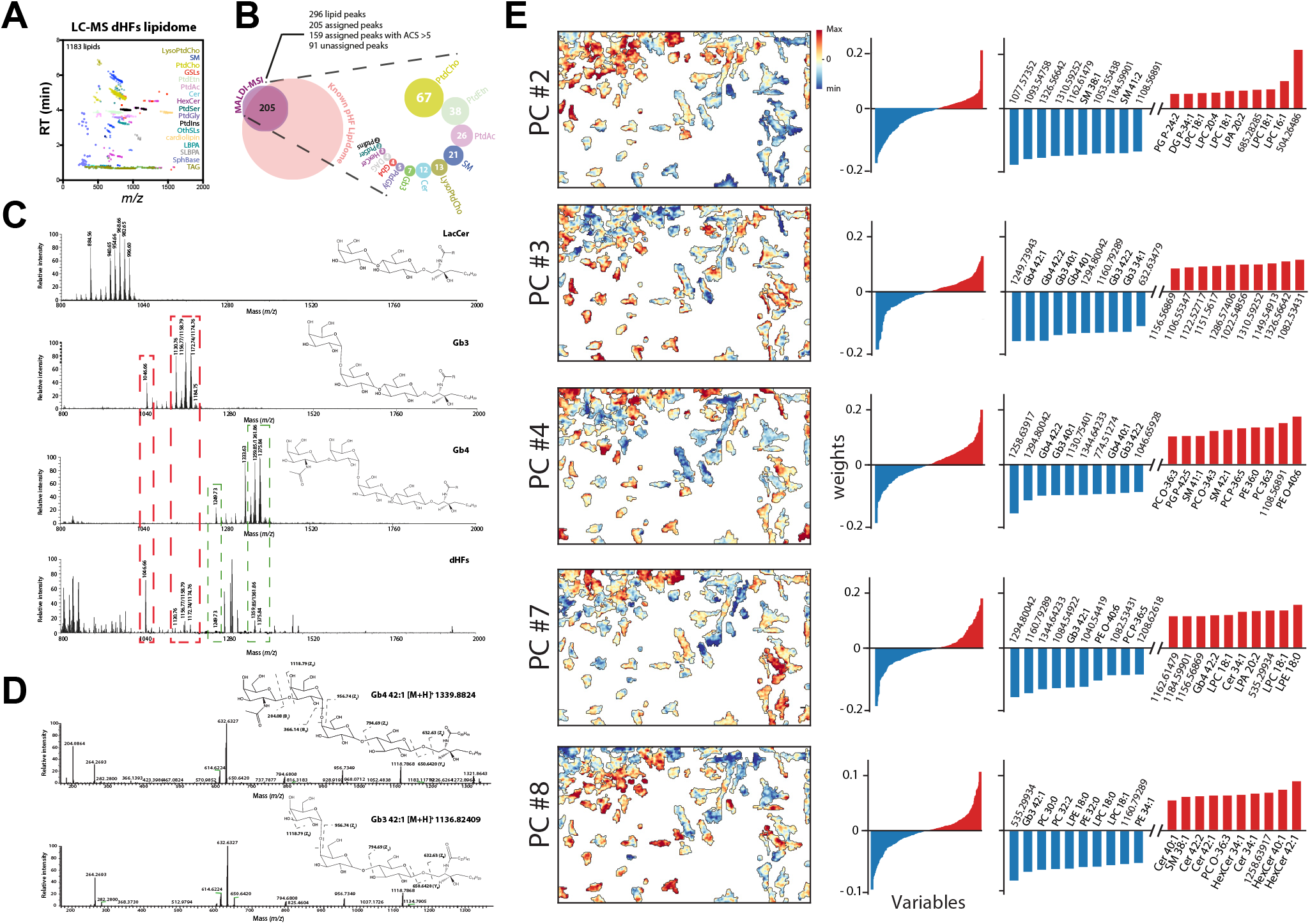
Lipid identification by standard lipidomics approaches and single-pixel analysis. (**A**) Scatterplot of the *m/z* values for the dHFs lipidome against their retention times as determined separating dHFs extracts on a HILIC column followed by ESI/MS positive-ion mode measurement.. Each dot represents a lipid and colour code is according to the lipid species they belong to. 1183 lipids were identified from 17 different lipid classes. (**B**) dHFs analysed by MALDI-MSI in positive-ion mode, returned 296 ion images in the lipids mass range belonging to 13 lipid classes as shown in the figure. By comparison with LC-ESI/MS, 205 lipids were identified, 159 of which with high confidence (ACS >5). (**C**) LactosylCeramide (LacCer), trihexosylceramide Gb3 and globosides Gb4 standards were analysed by MALDI-TOF in positive ion mode at 20 kV showing the presence of several species with different acyl chains. In the lower panel, MALDI-TOF mass spectra of lipids extracted from dHFs that were compared to the mass spectra of GSLs standards. (**D**) LC-MS/MS mass spectra of GSLs Gb3 and Gb4 (42:1) [M+H]^+^, from lipid extract. Precursor ion species detected at *m/z* 1339.8824 and at *m/z* 1136.8240 were isolated and MS/MS fragmentations were obtained upon HCD fragmentation at 23 eV. Corresponding fragmentation schemes identified the precursor ion species are shown. (**E**) The intensity values for each non-background pixel were used for principal component analysis. PC2, PC3, PC4, PC7 and PC8 values are displayed on the corresponding mass image. Positive (red) and negative (blue) contribution of variables to each principal component is shown. For every component, 10 loadings (lipids) that positively (red) or negatively (blue) contributed the most are shown.

**Figure S2.**
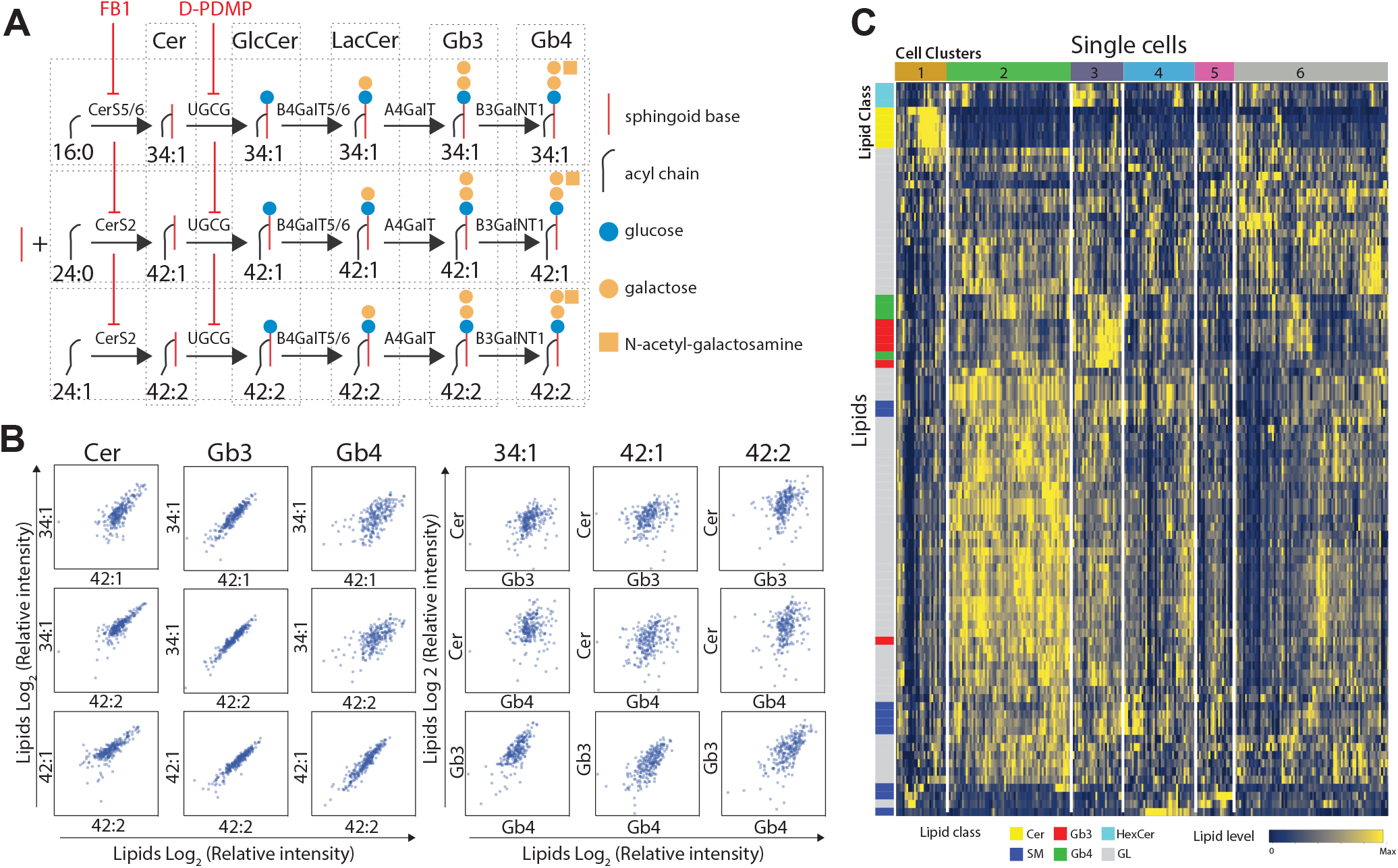
Analysis of lipid-lipid correlation. (**A**) Schematic depiction of the early sphingolipid synthetic pathway. *CerS*, ceramide synthases; *UGCG*, UDP-Glucose Ceramide Glucosyltransferase; *B4GalT5/6*, Beta-1,4-galactosyltransferase 5/6; *A4GalT*, Lactosylceramide 4-alpha-galactosyltransferase; *B3GalNT1*, Beta-1,3-N-acetylgalactosaminyltransferase 1. FB1 inhibits *CerS* and D-PDMP inhibits *UGCG*. (**B**) Batch corrected and normalized single-cell values of sphingolipids (i.e., Cers, and GSLs Gb3 and Gb4) are plotted as log2 relative intensities in each cell. Lipids with the same headgroup but different acyl chains (34:1, 42:1 and 42:2) are compared in the left panel. Lipids with different headgroups but the same acyl chain are compared in the right panel. (**C**) Heatmap of single cell lipidomics data. Columns and rows are ordered using optimal leaf ordering of the hierarchical similarity tree. Cells and lipids are annotated by cluster and class respectively. The 90 most variable lipids are displayed.

**Figure S3.**
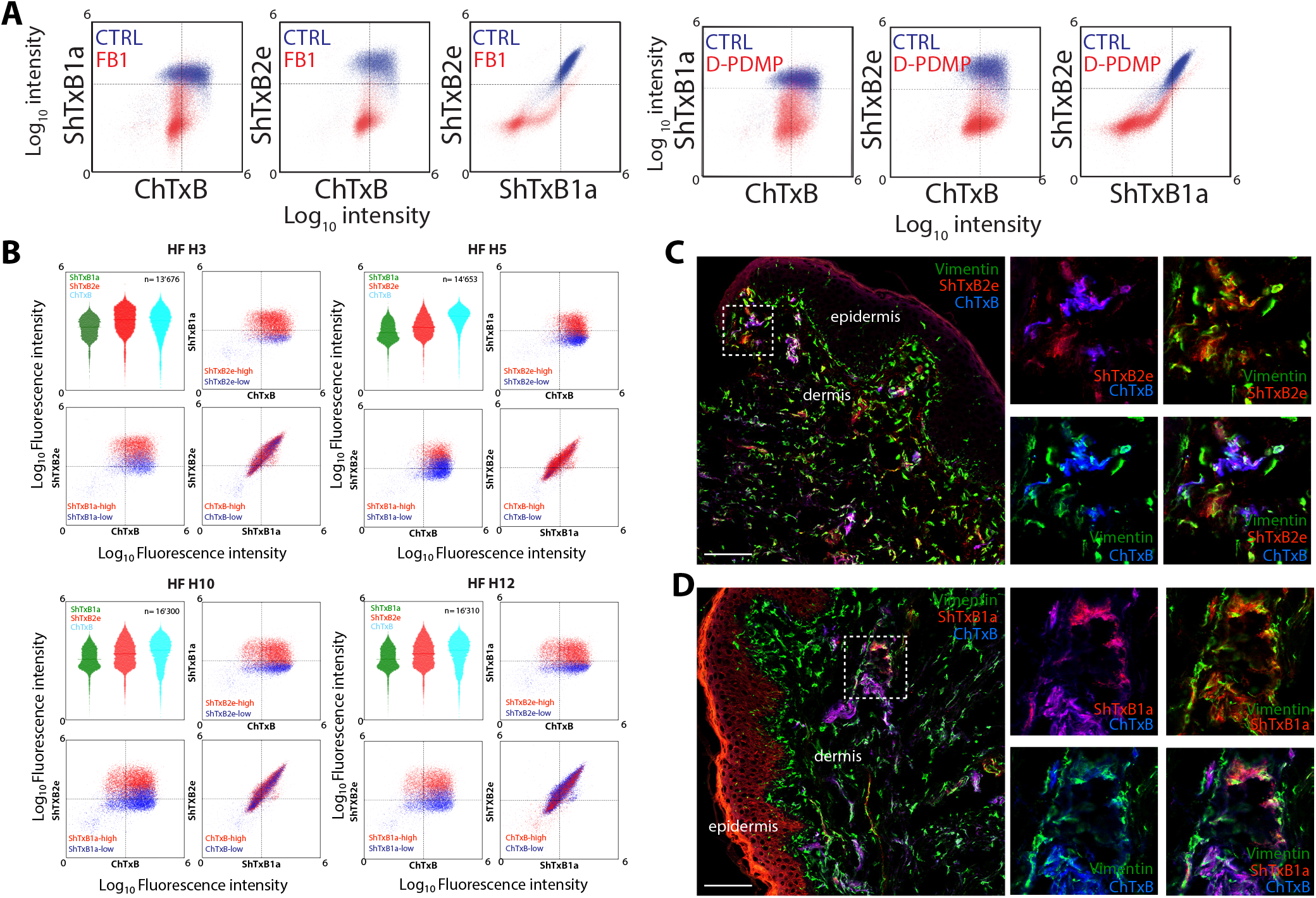
In vitro and in vivo assessment of lipid heterogeneity. (**A**) Cells were treated with SL inhibitors (25 μM FB1 or 10 μM D-PDMP), and stained with ShTxB1a, ShTxB2e and ChTxB. Scatter plots of fluorescence intensity values for each toxin are shown. Unstained cells were used as negative control and to determine the gates. CTRL cells are shown in blue; treated cells in red. CTRL, n= 23’806; FB1, n= 23’007; D-PDMP, n= 25’308. (**B**) Cytofluorometric analysis of four dHFs lines from healthy donors. Cells were stained with toxins recognizing lipids. Scatter plots and violin plots of fluorescence intensity values are shown for each toxin. Unstained cells were used as negative control and to determine the gates; n is indicated in the graph. (**C**,**D**) Confocal micrographs of human foreskin tissue sections, stained with bacterial toxins ShTxB1a/2e (red), and ChTxB (blue) and counterstained with antibodies against Vimentin (green). Scale bar is 100 μm.

**Figure S4.**
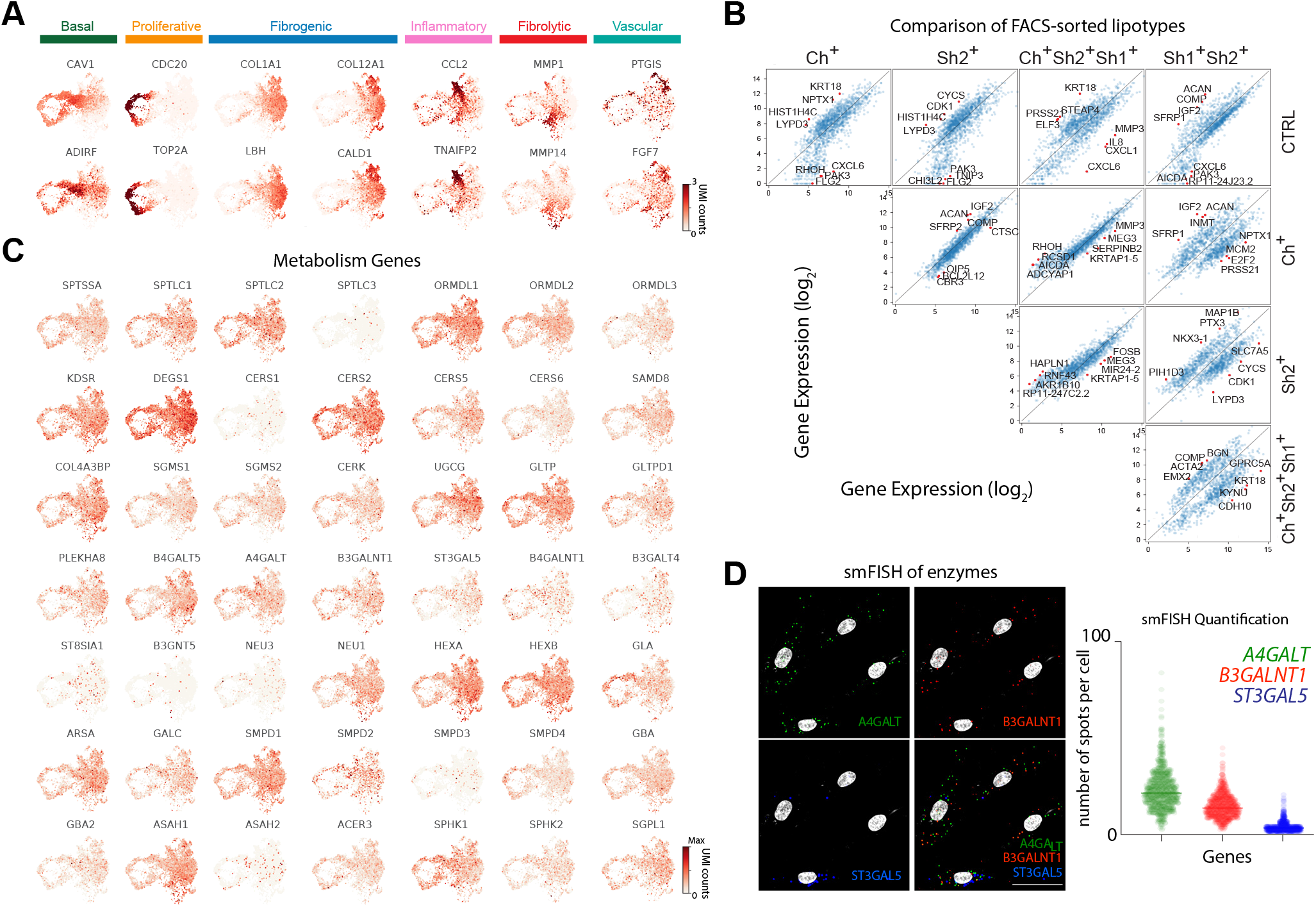
Gene expression profiles of dHF states and lipotypes. (**A**) UMAP embedding colored by the expression of markers for basal, proliferative, fibrogenic, inflammatory, fibrolytic and vascular cell states. (**B**) Scatterplot matrix comparing pariwise the gene expression of each lipotype. Names of the top differentially expressed genes are indicated for each comparison. (**C**) UMAP embedding colored by the expression of sphingolipid metabolic genes. (**D**) Representative images of mRNA-FISH staining using *A4GALT* (Gb3S), *B3GALN*T1 (Gb4S) and *ST3GAL5* (GM3S). Nuclei were labelled with Hoechst (grey). Scale bar is 50 μm. Right panel, the distribution of number of spots per cell for each gene as violin plots.

**Figure S5.**
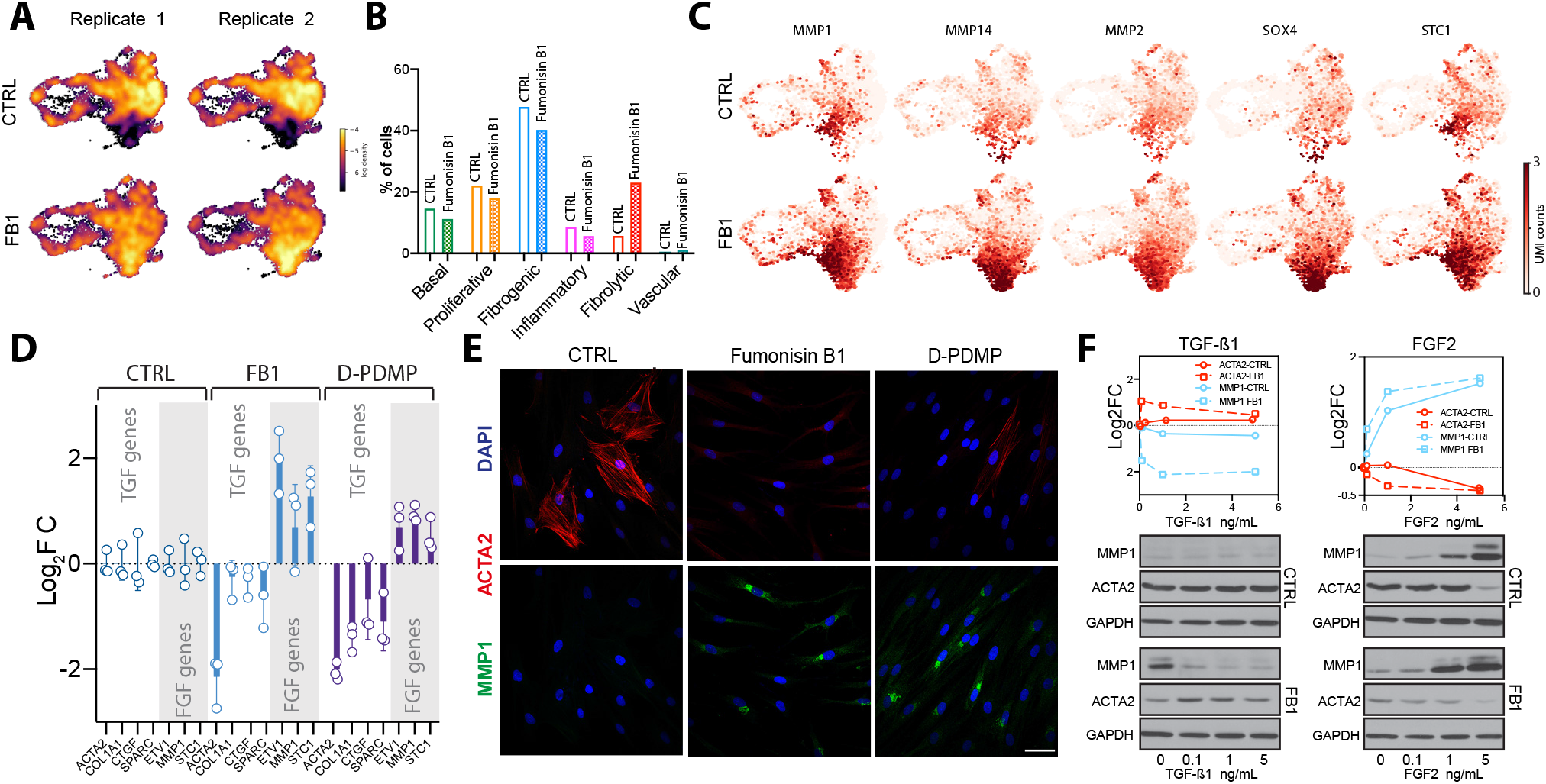
Effect of sphingolipid deprivation on dHFs transcription. (**A**) Density map of individual replicates of CTRL and FB1 treated cells in the UMAP space. (**B**) Quantification of cell distribution across basal, proliferative, fibrogenic, inflammatory, fibrolytic and vascular cell clusters in CTRL and FB1 treated cells expressed as percentage of total cells. (**C**) UMAP embedding colored by canonical markers for FGF2 signaling pathway in CTRL and FB1 treated samples. (**D**) Barplots of a qPCR quantifying the mRNA levels of TGF-β and FGF2 target genes in cells treated with 25 µM FB1 or 10 μM D-PDMP. qPCR data are shown as log2 fold changes over untreated cells (n=3; data are means ± StDev). (**E**) Confocal images of cells treated with 25 μM FB1 or 10 μM D-PDMP and stained with antibodies against ACTA2 (red) or MMP1 (green). Scale bar is 50 μm. (**F**) Plots indicate change in the relative amount of the indicated proteins (MMP1 and ACTA2) at increasing doses of FGF2 or TGF-β1 compared to untreated cells as determined by densitometry. Cells were treated with 25 μM FB1, serum starved for 8h and treated with increasing concentration of FGF2 or TGF-β1 for 72h. Cells were then lysed and processed for SDS-PAGE and western blotting analysis with antibodies recognizing ACTA2 and MMP1. Quantification is shown in the upper panels. Data were normalized against GAPDH and shown as log2 fold changes over control; n=4.

**Figure S6.**
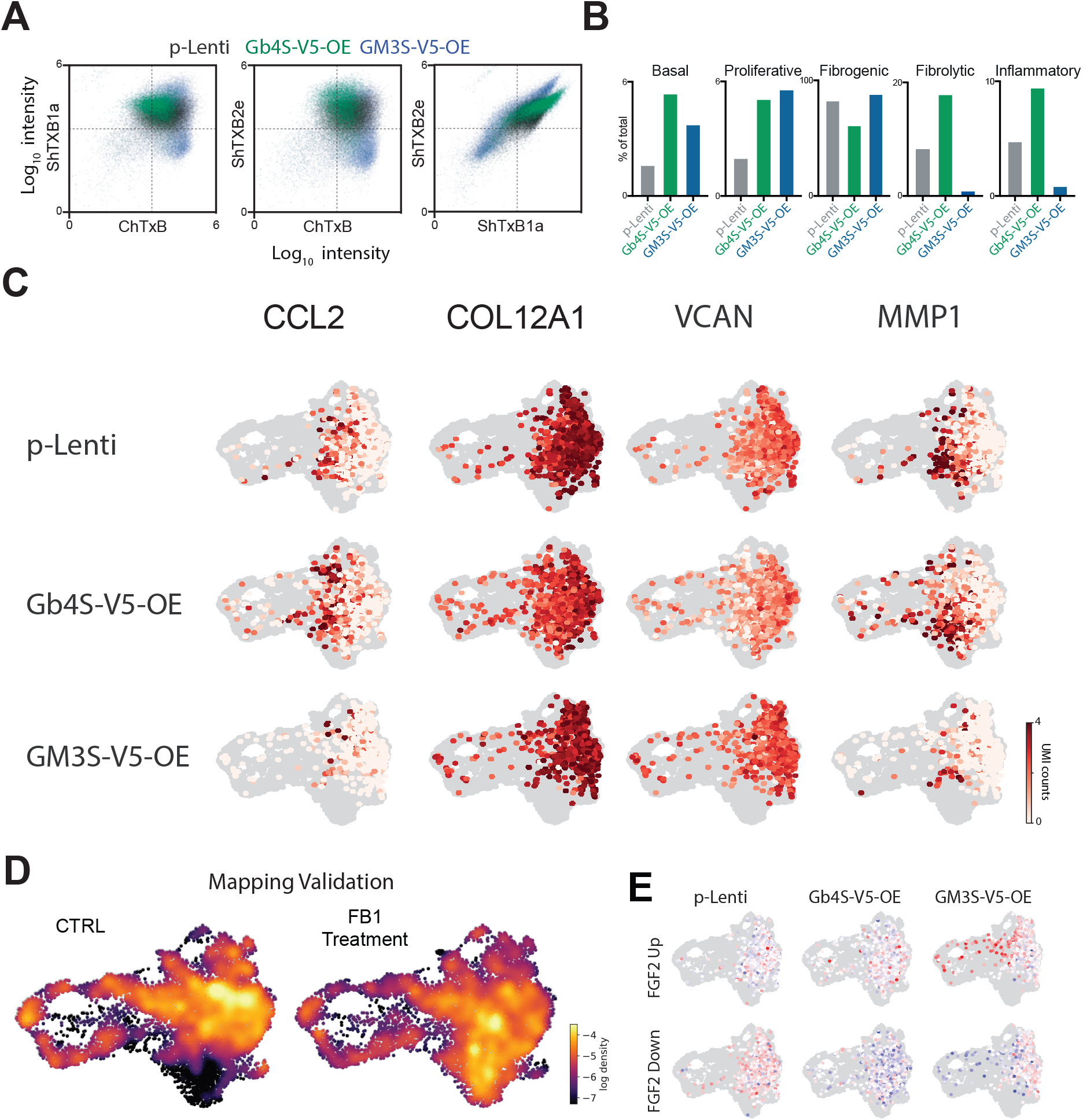
Cytofluorimetric and gene expression evaluation of the effects of sphingolipid manipulations. (**A**) Cytofluorometric analysis with bacterial toxins. Cells were stained with toxins. Scatter plots of fluorescence intensity values are shown for each toxin. Unstained cells were used as negative control. Cells were gated on the singlet population; CTRL n=23’858; GM3S-OE n=24’007; Gb4S-OE n=11’358. (**B**) Quantification of cell distribution across basal, proliferative, fibrogenic, inflammatory and fibrolytic cell clusters in pLenti, Gb4S-OE and GM3S-OE cells expressed as percentage of total cells. (**C**) UMAP embedding colored by the gene expression levels canonical markers for FGF2 or TGF-β signaling pathways. (**D**) Single-cell transcriptomes from the Control and FB1 treated dHF samples mapped onto the embedding of Fig. 5A (*i*.*e*. their own) to validate the similarity-based mapping procedure used to produce Fig. 5G. (**E**) FGF2 gene signatures for overexpression lines and control cells overlayed onto the UMAP embedding.

## SUPPLEMENTAL TABLEs

**Table S1. Putative lipid features (296 total) recorded by AP-SMALDI10 and AP-SMALDI5 AF from dHFs in positive ion mode with their experimental** *m/z* **values**. Lipid features confirmed by LC/MS and MRM are annotated. Data refer to four separate MSI measurements conducted on independently prepared samples. Analysis were obtained with mass resolving powers of 240,000 (@*m/z* 200). Lipids were annotated by the workflow described in the methods.

**Table S2. List of chemicals used in this study**.

**Table S3. List of vectors used for lentiviral transduction of dHFs**.

**Table S4. List of the antibodies used in this study for flow cytometry (FACS), Immunofluorescence (IF) and Western Blotting (WB)**.

**Table S5. List of Real-time primers used in this work**.

## METHODS

### CONTACT FOR REAGENTS AND RESOURCE SHARING

Further information and requests for resources and reagents should be directed to and will be fulfilled by the Lead Contacts: Giovanni D’Angelo and Gioele La Manno

#### Cell lines and cell growth conditions

Dermal human fibroblasts (dHFs) and primary foreskin dHF expressing dominant-negative FGFR1 (DNFGFR1) or dominant-negative TGFBR2 (DNTGFBR2) were kindly provided by the laboratory of Prof. Paolo Dotto (DB-UNIL, Lausanne, Switzerland). Cells were grown in DMEM supplemented with 10% (v/v) foetal bovine serum (FBS), 4.5 g/l glucose, 2 mM L-glutamine, 1 U/ml penicillin/streptomycin. The cell lines were grown under a controlled atmosphere in the presence of 5% CO_2_ at 37°C. Cells were grown in a flask until 90% confluence. The medium was removed and trypsin-EDTA solution (0.05% trypsin, 0.02% EDTA) was added for 3-5 min. Then, the medium was added back to block the protease action, and the cells were collected into a plastic tube. After centrifugation for 5 min at 1200 rpm, the cell pellet was resuspended in fresh medium, gently mixed, and placed in a new plastic flask with fresh growth medium.

#### Generation of stable cell line over-expressing GSLs synthesizing enzymes

For lentiviral production 293T cells were transfected in 10 cm dishes with 4 µg of the corresponding lentiviral vectors (empty vector or transgene listed in Table S3) and packaging vectors pMD2.G (1 µg) and pCMVR8.74 (3 µg) using Lipofectamine 2000 (1:400 in OPTIMEM). The next day, the medium was changed for DMEM with 10% FCS and penicillin/streptomycin. The first harvest of viral particles was done 40 hours after transfection and fresh medium was added to the cells. Virus-containing medium was filtered through 0.45 µm filters and 10ml of filtrate was applied to 1 million cells in 15 cm dishes. The second harvest was done 24 hours after the first harvest and the transduction repeated as described. The next day, transduced cells were split and selected with Blasticidin (10 µg/mL) for at least one week.

#### MALDI-MSI workflow -Sample preparation

dHFs, were directly grown on glass bottom culture dishes (Mattek) in complete media to roughly 60% confluence. After aspiration of media, cells were washed twice with PBS, followed by fixation in 0.25% glutaraldehyde for 15 minutes. Following fixation and washing, cells were stained, when required, with fluorescent dyes or with bacterial toxins (as described below). Pen marks were manually drawn on the glass slide, on the back side of the sample, for image registration and then, confocal images (as described below) were acquired from an area of interest. For MALDI-MSI analyses, 150 μL of 2,5-dihydroxybenzoic acid (DHB), (30 mg/mL in 50:50 acetonitrile/ water/0.1% TFA), were deposited on the surface of the samples using the automatic SMALDIPrep (TransMIT GmbH, Giessen, Germany).

#### MALDI-MSI workflow -Analysis

MSI experiments were performed using the AP-SMALDI10 or AP-SMALDI5 AF systems that couple a Q Exactive orbital trapping mass spectrometer (Thermo Fisher Scientific, Bremen, Germany) with an atmospheric-pressure scanning-microprobe MALDI imaging source (AP-SMALDI, TransMIT GmbH, Giessen, Germany). The MALDI laser focus was optimized manually using the source cameras aiming at a diameter of the focused beam of 5 μm. For each pixel, the spectrum was accumulated from 50 laser shots at 100 Hz. MS parameters in the Tune software (Thermo Fisher Scientific) were set to the spray voltage of 4 kV, S-Lens 100 eV, capillary temperature to 250 °C. The step size of the sample stage was set to 7 or 5 mm. Positive ion mode measurements were performed in full scan mode in the mass range *m/z* 400-1600 with a resolving power set to R = 240000 at *m/z* = 200. Mass spectra were internally calibrated using the lock mass feature of the instrument.

#### MALDI-MSI images generation and annotation

The data obtained were converted from the RAW format into the imzML format containing only centroided data using the RAW2IMZML software.

Mass images (296) were generated by MSiReader software [58] after TIC normalization with a *m/z* tolerance of 3 ppm. Metabolite annotation was performed in two steps: annotation with public databases and software and then ESI-LC/MS and MRM confirmation (as described below).

imzML files were uploaded on METASPACE [59] for a preliminary lipids annotation, with the *m/z* tolerance of 3 ppm and FDR of 5-50% against the Swiss Lipids metabolite database.

For further confirmation, RAW data were analyzed with the software Alex123 (Analysis of Lipids Experiments)[60] using a library with curated lipid ionization information with a *m/z* mass tolerance of 0.005.

After peaks identification based on the combination of two step approaches, an attribution confidence score (ACS) was associated to each lipid as detailed: 1 point was attributed when a lipid was identified by Alex123 software, while 1,2,3 or 4 points were attributed when a lipid was identified on MetaSpace with FDR 50, 20, 10 or 5% respectively. Finally, 5 points were given to a lipid that was identified by ESI-LC/MS with 1 extra point if the lipid identity was confirmed by MRM. Only lipids with an ACS > 5 were considered identified with high confidence.

#### Single pixel analysis

Single-pixel analysis was performed using Fiji software[61] on the 296 mass images collected at 7 μm spatial resolution by AP-SMALDI10 (TransMit GmbH, Giessen, Germany) and reconstructed by Mirion software [62]. Pixels corresponding to area where no cells were present displayed an extremely low total lipid intensity. These areas were identified using the evenly distributed lipid PtdAc 38:4 (*m/z* = 747.4937) and removed from further analysis. Lipid intensities in each pixel belonging to cell areas were then log-transformed and standardized across the entire image. PCA analysis was performed and the coordinate of each component was displayed with a different colour. The absolute values of the PCA loadings were then used to identify the lipids with most of the variance of each single component. The first level clustering was performed on 12 principal components (PCs) with the Leiden algorithm (in leidenalg implementation) using default parameters..

#### Single-cell lipids data analysis

Single dHFs were manually segmented on MsiReader with the free hand tool for ROI selection, after TIC normalization. Raw abundance data for each scan and each pixel in a ROI were exported with the MSiExport tool for the 296 *m/z* in the lipid mass range.

Normalized lipid count values for biological replicates were integrated using ComBat [57] to correct batch effects. Batch corrected counts per each lipid were then scaled by their average. The coefficient of variation (CV) was computed for each lipid using the normalized values. Lipid co-variation was evaluated using the Pearson Correlation Coefficient (Pearson’s R) among cells.. Lipids with R >0.85 were connected with edges to build a lipid correlation network using Cytoscape 3.8.0. Node size was calculated using the CVs (bigger is the node bigger is the CV and vice versa) while node transparency was calculated according to the ACS (as detailed above).

Similarly, single-cell lipidome values were used to determine their co-variation using the Pearson correlation R. Cells with R >0.35 were connected to build a cell correlation network using Cytoscape 3.8.0. Clustering was performed using the ClusterViz App of Cytoscape 3.8.0 and according to the fast agglomerate algorithm FAG-EC.

#### Lipid extraction

Total lipid extracts were prepared using a standard MTBE protocol followed by a methylamine treatment for total lipid analysis by mass spectrometry [63]. Briefly, cell pellet was resuspended in 100 μL H_2_O. 360 μL methanol and 1.2 mL of MTBE were added and samples were placed for 10 min on a vortex at 4 °C followed by incubation for 1 h at room temperature on a shaker. Phase separation was induced by addition of 200 μL of H_2_O. After 10 min at room temperature, samples were centrifuged at 1000 g for 10 min. The upper (organic) phase was transferred into a glass tube and the lower phase was re-extracted with 400 μL artificial upper phase [MTBE/methanol/H_2_O (10:3:1.5, v/v/v)]. The combined organic phases were dried in a vacuum concentrator. Lipids were then resuspended in 500 μL of CHCl_3_ and divided in two aliquots for a further methylamine treatment for sphingo-and glycosphingolipids analysis. In details, 500 μL of freshly prepared monomethylamine reagent [methylamine/ H_2_O/n-butanol/methanol (5:3:1:4, (v/v/v/v)] was added to the dried lipid extract and then incubated at 53 °C for 1 h in a water bath. Lipids were cooled to room temperature and then dried. The dried lipid extract was then extracted by n-butanol extraction using 300 μL water-saturated n-butanol and 150 μL H_2_O. The organic phase was collected, and the aqueous phase was re-extracted twice with 300 μL water-saturated n-butanol. The organic phases were pooled and dried in a vacuum concentrator.

#### MALDI-TOF untargeted lipidomics

Extracted lipids were resuspended in 500 μL of CHCl_3_ and analyzed by MALDI-MS. 30 mg/mL 2,5-DHB was freshly prepared in acetonitrile/water solution (50:50 v/v) with 0.1% TFA. An equivalent volume of sample solution (50 μL) was then mixed with matrix before deposition on the MALDI target. All mass spectrometry analysis for the identification of lipids (*m/z* 400-1800) were obtained using an Applied Biosystems 4800 MALDI-TOF/TOF mass spectrometer equipped with a 200 Hz tripled-frequency Nd:YAG pulsed laser with 355 nm wavelength. Measurements were performed in positive ion reflection mode at an accelerating potential of 20 kV. Each mass spectra were obtained by applying a laser energy of 4600 watts/cm^2^, averaging 4000 single laser shots/spectrum.

#### LC-MS untargeted lipidomics

For phospholipid analysis, lipid extracts (2 μL injection volume in CHCl_3_:MeOH 2:1) were separated over an 8 minute gradient at a flow rate of 200 μL/min on a HILIC Kinetex Column (2.6lm, 2.1 × 50 mm^2^) on a Shimadzu Prominence UFPLC xr system (Tokyo, Japan). Mobile phase A was acetonitrile:methanol 10:1 (v/v) containing 10 mM ammonium formate and 0.5% formic acid while mobile phase B was deionized water containing 10 mM ammonium formate and 0.5% formic acid. The elution of the gradient began with 5% B at a 200 μL/min flow and increased linearly to 50% B over 7 min, then the elution continued at 50% B for 1.5 min and finally, the column was re-equilibrated for 2.5 min. MS data were acquired in full-scan mode at high resolution on a hybrid Orbitrap Elite (Thermo Fisher Scientific, Bremen, Germany). The system was operated at 240,000 resolution (*m/z* 400) with an AGC set at 1.0E6 and one microscan set at 10-ms maximum injection time. The heated electrospray source HESI II was operated in positive mode at a temperature of 90 C and a source voltage at 4.0KV. Sheath gas and auxiliary gas were set at 20 and 5 arbitrary units, respectively, while the transfer capillary temperature was set to 275 °C.

Mass spectrometry data were acquired with LTQ Tune-plus2.7SP2 and treated with Xcalibur 4.0QF2 (Thermo Fisher Scientific). Lipid identification was carried out with Lipid Data Analyzer II (LDA v. 2.6.3, IGB-TUG Graz University) [64]. The LDA algorithm identifies peaks by their respective retention time, *m/z* and intensity. Care was taken to calibrate the instrument regularly to ensure a mass accuracy consistently lower than 3 ppm thereby leaving only few theoretical possibilities for elemental assignment.

Data visualization was improved with LCMSexplorer in a homemade web tool hosted at EPFL.

MS/MS spectrum were acquired using a hybrid Velos pro dual cell differential pressure linear ion trap mass spectrometer with a high field Orbitrap Elite instrument. The fragment spectra were generated using parallel MS and MSn analysis with 23eV in the HCD cell.

#### TSQ ANALYSIS (Multi Reaction Monitoring)

The lipid extracts were normalized against total phosphate content before analysis. Dried samples were re-suspended in 100uL chloroform/methanol (1:2) containing 5mM ammonium acetate and transferred to a 96well plate.

The sphingolipids species were quantified using multiple reaction monitoring MRM on a TSQ vantage Extended mass range Mass spectrometer (Thermo Fisher Scientific) equipped with a robotic nanoflow ion source (Triversa Nanomate, Advion Biosciences). Auto-tuned collision energies and s-lens values were used on standards lipids covering the analyzed subclasses to improve MRM transitions. Mass spectrometry data were acquired with TSQ Tune 2.6 SP1 and Xcalibur 4.0QF2 (Thermo Fisher Scientific).

The mass spectrometry results were treated with an automatic MRM lipid quantification platform, developed and hosted at EPFL Lausanne Switzerland (http://lipidomes.epfl.ch). Areas under the curve of MRM transitions were quantified relative to the internal standard.

#### HPTLC

dHFs were pulse-labelled in serum-free DMEM, supplemented with 1% BSA fatty acid free, with 0.1 mCi/mL H^3^-sphingosine for 2 h. After labelling, the cells were further incubated in DMEM + 10% FCS for 24 h. Cells were then harvested and lipids extracted with the standard Bligh and Dyer protocol (Bligh EG and Dyer WJ, 1959). Dried lipids were resuspended in 150 ml of CHCl_3_ and spotted on silica gel high-performance–TLC (HPTLC) plates (Merck, Germany) and resolved with a mixture of chloroform, methanol and water (65:25:4 v/v/v). To visualize and analyse radiolabelled sphingolipids (i.e. Cer, GlcCer, LacCer Gb3, GM3 and SM), the TLC plates were placed in the RITA TLC Analyser (Raytest, Germany) and quantified using GINA (Raytest, Germany) software analysis. The percentage of total C.P.M. associated with Cer, GlcCer, LacCer, Gb3, GM3 and SM peaks for each of the lipids is reported.

#### Immunofluorescence analysis

For IF analysis, dHFs were grown to approximately 80% confluency on glass coverslips, fixed with 4% paraformaldehyde for 10 minutes at RT and then washed three times with PBS. For tissues, samples were embedded in OCT compound and frozen, to be subsequently cryosectioned with a cryotome into 7-8 μm sections. Before staining, sections were dried at room temperature and then fixed with 4% paraformaldehyde for 20 minutes. After fixation, cells and tissues were blocked with 5% BSA and permeabilized when required with 0.5% saponin for 20 minutes at RT, followed by a 1 h incubation with selected antibodies (listed in Table S4) against the antigen of interest in blocking reagent. Cells and tissues were then washed three times with PBS and incubated with appropriate isotype-matched, AlexaFluor-conjugated secondary antibodies (Invitrogen, USA) diluted in blocking solution for 30 min. After immuno-staining, they were washed three times in PBS and once in water, to remove salts. After Hoechst staining for nuclei, the samples were mounted with Fluoromount-G and analyzed under a confocal microscope Zeiss LSM700 with 20x air objective (0.8 NA) or Leica SP8 with 10x air objective (0.3 NA) or 20x air objective (0.75 NA).

Optical confocal sections were taken at 1 Airy unit under non-saturated conditions with a resolution of 1024×1024 or 2048×2048 pixels and frame average 4. Images were then processed using Fiji software [61]. Adobe Photoshop CS3 and Fiji softwares were used to adjust the contrast of the images, whereas Adobe Illustrator 2020 was used to illustrate figures.

#### Fluorescence staining with toxins

For ShTxB1a-Cy3 and ChTxB-AlexaFluor488 or ChTxB-AlexaFluor647 staining, the cells were fixed with 4% PFA, blocked in PBS containing 5% bovine serum albumin (BSA) without detergent, incubated with fluorescently labelled B-subunit toxins for 1h, and then mounted with Fluoromount-G. In the case of ShTxB2e, after fixation and blocking, cells were incubated with the bacterial toxin for 30 min at RT, followed by conjugation with primary antibody for 1 hour and fluorescently-labelled secondary antibodies for 30 minutes. Cells were analysed by confocal microscopy as described above.

#### Correlative toxin staining/MALDI-MSI analysis

Cells were seeded the day before on a gridded glass bottom culture dish (Mattek) in complete media to reach the day after roughly 70% confluence. Cells were then fixed and stained with toxins as described above and analyzed by confocal microscopy and phase contrast to recognize the grids for subsequent registration. Images were acquired with a confocal microscope Leica SP8 with 20x air objective (0.75 NA). Optical confocal sections were taken at 1 Airy unit under non-saturated conditions with a resolution of 1024×1024 pixels and frame average 4 with the LAS X navigator tool to obtain tiles. Marks were drawn on the glass slide to assist image registration and samples were processed for MALDI-MSI analysis as described above.

#### Flow cytometry analysis

dHFs were subjected to trypsin digestion (0.05% Trypsin/ EDTA) and washed twice with PBS. For cell-surface GSLs staining, resuspended cells were blocked with 2% BSA for 30 minutes at 4 °C. Then, cells were extensively washed with PBS and incubated with optimal concentrations of the toxins for 1 h at 4 °C and after washing with primary antibody. Cells were then washed again and incubated with fluorescently-labelled secondary antibodies when required for 30 minutes. Cells resuspended in 2% BSA were analyzed by BD LSR Fortessa or LSRII SORP (Becton Dickinson). Unlabeled cells were used as negative control. Viable cells were gated, and GSLs expressions were further analyzed in the gated region. Antibodies used are described in Table S4. Data was exported with an in-house built R script and analysed with GraphPad Prism 8 software.

#### Isolation of lipotypes by FACS

A triplicate of primary human fibroblasts (5×10^6^) in suspension were collected and stained as described above. Cells were resuspended in 2% BSA and sorted using a flow cytometer (FACSAriaII). After gating, four populations (ChTxB^+^, ShTxB1a/ShTxB2e, triple positive and ShTxB2e^+^) were directly sorted through a 100 mm nozzle at 4 °C in 5 mL tubes filled with 1 mL lysis buffer or complete media. Cells were sorted in continuous to get the maximum amount from each population. Unlabeled cells were used as negative control.

#### Drug treatments

dHFs were treated by adding inhibitors of SLs synthesis, Fumonisin B1 25 μM, D-PDMP 10 μM or Myriocin 2.5 μM, in complete media for six days. Stock solutions of the drugs were prepared dissolving powders in DMSO following providers instructions.

#### FGF2/TGF-β1 treatment

dHFs were serum starved for 24h and then treated with different concentration (0, 0.1, 1 or 5 ng/mL) of growth factors FGF2 and TGF-β1 by adding them in complete media for 72 hours. TGF-β1 stock solution at 50 mg/mL was prepared by dissolving it in water with citric acid 10mM pH3 and 0.1% BSA. FGF2 stock solution at 10 mg/ mL in PBS with 0.5% BSA.

#### Fluorescence In Situ Hybridization (FISH)

RNAscope Multiplex Fluorescent V2 assay (Bio-techne, Cat. No. 323110) was performed according to manufacturer’s protocol [66] on cells cultured in chamber slides, hybridized with the probes Hs 3plex positive control (Biotechne, Cat. No. 320861) or 3Plex negative control (Bio-techne, Cat. No. 320751) or Hs-*A4GALT*-C1 (Bio-techne, Cat. No. 486601), Hs-*ST3GAL5*-C2 (Bio-techne, Cat. No. 816191-C2) and Hs-*B3GALN*T1-C3 (Bio-techne, Cat. No. 816181–C3) simultaneously at 40°C for 2 hours. The different channels were revealed with TSA Opal520 (Akoya Biosciences, Cat. No. FP-1487001KT) for C1, TSA Opal650 (Akoya Biosciences, Cat. No. FP1488001KT) for C2 and TSA Opal570 (Akoya Biosciences, Cat. No. FP1488001KT) for C3. Cells were counterstained with DAPI and mounted with Prolong Diamond Antifade Mountant (Thermo Fisher, P36965).

Images were acquired with a confocal microscope Zeiss LSM700 with 20x air objective (0.8 NA). Optical confocal sections were taken at 1 Airy unit under non-saturated conditions with a resolution of 1024×1024 pixels and frame average 4. The acquired images were analyzed with Fiji [61] using a custom ImageJ script. Briefly, the nuclear staining was used to create a nuclei mask using the Huang automatic thresholding method and cleaned for “dust”/”noisy pixels” using a median filter (radius 5 pixel). The median filter radius was selected based on average nuclei length (50 pixels). This nuclei mask was used to generate a cell mask via the Voronoi operation, 3) the cell mask was used to create Regions Of Interest (ROIs) of the cells, 4) for each FISH staining channel and for each cell ROI, the FISH spots were detected as local maxima. The script outputs a result table with the count of FISH spots detected per channel, and a tiff file with the cell ROIs and the FISH spots ROIs overlaid.

#### Correlative fluorescence/mRNA-FISH analysis

Cells were seeded the day before on a gridded glass bottom culture dish (Mattek) in complete media to reach the day after roughly 70% confluence. Cells were then fixed and stained with fluorescently labelled toxins ShTx-B1a-Cy3 and ChTxB-488 as described above and analyzed by confocal microscopy and phase contrast to recognize the grids for subsequent registration.

Images were acquired with a confocal microscope Zeiss LSM700 with 20x air objective (0.8 NA). Optical confocal sections were taken at 1 Airy unit under non-saturated conditions with a resolution of 1024×1024 pixels and frame average 4. Cells were kept for the entire staining procedure in RNAlater (Sigma) to stabilize RNA molecules before FISH staining. After confocal imaging, cells were then processed for mRNA-FISH with the probes Hs-*A4GALT*-C1 (Bio-techne, Cat. No. 486601) and Hs-*ST-3GAL5*-C2 (Bio-techne, Cat. No. 816191-C2) as described above. Images were acquired with the same microscope used after toxin staining with a resolution of 2048×2048 pixels. Correlative images were mounted manually with the Fiji software with the help of the phase contrast images of the grids. mRNA-FISH images were analyzed using a custom ImageJ script as described above.

#### Bulk RNA-sequencing of FACS sorted dHFs population

Bulk RNA sequencing was performed on the following FACS sorted populations of dHFs: Cholera Toxin positive (n=2), ChTxB/ShTxB1a/ShTxB2e positive (n=2), ShTxB1a/ShTxB2e positive (n=3) and ShTxB2e positive (n=1) and to a control unsorted population (n=2). Total RNA was isolated from FACS sorted dHFs populations using Rneasy Mini kits (Qiagen, Germany) according to the manufacturer instructions. The yield and the integrity of the RNA were determined using a spectrophotometer (NanoDrop ND-1000; Thermo Scientific, USA).

Total RNA (10ng-1mg, depending on the different population) were submitted for RNA-seq with GENEWIZ, NJ.

Libraries were prepared using Illumina HiSeq platform with ultra-low input configuration and sequenced with 2× 150 bp sequencing configuration to a depth of 350 million reads (GENEWIZ, NJ).

Sequence reads were trimmed to remove possible adapter sequences and nucleotides with poor quality using Trimmomatic v.0.36. The trimmed reads were mapped to the Homo sapiens GRCh38 reference genome available on ENSEMBL using the STAR aligner v.2.5.2b.

Unique gene hit counts were calculated by using featureCounts from the Subread package v.1.5.2. The hit counts were summarized and reported using the gene_id feature in the annotation file. Only unique reads that fell within exon regions were counted (GENEWIZ, NJ). These bulk quantifications were used to extract, for each of the subpopulations, a set of enriched genes. First, we filtered the data tables from the genes that were lowly expressed along most of the samples: genes detected at average depth of less than 1.5 reads and that were at 8 reads in at least 15% of the samples were discarded. Samples were depth-normalized to reads per million (RPM) and log transformed. Then the average level of expression of a gene each condition was compared with the average of the rest of the samples. Genes were, then, sorted by the fold increase, excluding genes for which log2RPM was less than 4. For each of the subpopulations the top 200 genes were used to compute the population signature on the single-cell data (described below).

The signatures for TGF-β and FGF2 were computed with the same procedure starting from a list of the top 200 genes extracted from a previous bulk RNA-seq experiment [40].

#### scRNAseq experiment

scRNAseq experiments were performed using 10X Genomics Chromium scRNAseq kit v3.1. 3500 cells were loaded for each of the reaction following provider instructions. Libraries were sequenced at the depth of 300 million reads corresponding to an average of 80k reads per cell. Data was pre-processed using cellranger and velocyto v0.17.

#### scRNA-seq experiments -WT and FB1 treatment

We selected most variable genes using a CV-mean modeling-based feature selection with minimal dispersion 0.5, maximal mean 3 and minimal mean 0.0125 as previously described [67]. Single-cell profiles were normalized by the total UMI count andlog-normalized.PCA was performed retaining the top 50 of components.

The cluster analysis and embedding of Control and FB1 treatment datasets were performed on the integrated dataset. Gene expression visualization and differential expression analysis were performed using the data before integration. The integration was performed using the Seurat integration algorithm. Clustering was performed using the Louvain clustering algorithm, with default parameters. Clusters were annotated consulting the literature relative to the differential expressed features.

Density estimation on the embedding was performed using a kernel density estimator (scikit-learn implementation). Signature enrichment scores for the toxin-marked subpopulations and for TGF-β and FGF2 pathways were both computed as average Z-scores across the 200-genes lists described above. We used a procedure analogous to the one used Seurat CellCycleScoring function [68]. Briefly, the log-transformed depth-corrected counts of genes in the list were zero-centered and standardized and the obtained Z-scores were averaged along each cell. To avoid capturing sequencing depth bias the score was corrected for the expectation estimated computing the average Z score of a random sample of genes stratified by average expression level.

#### scRNA-seq -Cell lines over-expressing GSLs synthesizing enzymes

Single-cell data was preprocessed excluding from cells with low sequencing depth (counts less than 1/3rd of the sample median). Then a CV-mean modeling-based feature selection procedure was performed to select the 500 most highly overdispersed genes. To adjust for different sequencing depths, UMI counts were downsampled (using the downsample_counts scanpy function) by cell-specific factors computed so that the median count distribution of each sample matched the one of the sample with lowest median.

To map single-cell gene expression profiles onto the same embedding computed on WT and FB1-treated cells we used a similarity based placement approach. Specifically, we assigned to each cell the UMAP coordinates corresponding to the median of the 20 most similar WT-FB1 cells in gene expression space. Similarity was computed as inverse of the euclidean distance in the log transformed data.

To compute the TGF-β and FGF2 pathways signature scores we used the same genes and procedure as described above. To make both the density plots and signature scores more directly comparable we have performed the above analysis randomly downsampling the cells to 800 per replicate.

#### Real Time PCR

Total RNA was extracted and DNAse treated from a 10-cm dish with RNeasy Kit (Qiagen, Germany), according to manufacture instruction. The yield and the integrity of the RNA were determined using a spectrophotometer (NanoDrop ND-1000; Thermo Scientific, USA). Reverse transcription was performed using 250ng of RNA using random primers and SuperScript II (Invitrogen). Real-time PCR was performed with 7900HT Fast Real-Time PCR système (Applied Biosystems) using PowerUp SYBR Green reagent for detection (Applied Biosystems). All primer sequences are listed in Table S5. mRNA levels were normalized to three housekeeping genes: Hypoxanthine-guanine phosphoribosyltransferase (HPRT), b-microglobulin (bM2), TATA-binding protein (TBP). Data were average of three replicates from independent experiments.

#### SDS-PAGE and western blotting

For sample preparation, after treatment, the cells were washed three times with PBS and lysed in RIPA buffer (150 mM NaCl, 1% Triton X-100, 0.5% sodium deoxycholate, 0.1% SDS, 25 mM Tris-HCl, pH 7.4), supplemented with protease cocktail inhibitor. The lysates were clarified by centrifugation, and quantified using a commercially available BCA kit (Pierce^™^ BCA Protein Assay Kit, ThermoFisher) according to the manufacturer instructions. Samples were prepared by adding an equal volume of 2x SDS sample buffer, denaturated at 95 °C for 5 min and resolved by SDS-PAGE and immunoblot.

For immunoblotting, the strips containing the proteins of interest were blocked in TBS-T/ 5% BSA for 45 min at RT, and then with the primary antibody diluted at its working concentration in the blocking solution buffer overnight at 4 °C. After Washing with TBS-T, the strips were next incubated for 1 h with the appropriate HRP-conjugated secondary antibody, diluted in antibody dilution buffer and washed twice in TTBS, for 10 min each. After washing, the strips were incubated with the ECL solution for 3 minutes and exposed to x-ray films, which were then scanned. The intensity of the bands and preparation of images was done using ImageJ and Adobe Illustrator 2020.

#### Statistical evaluation

Statistical testing was performed using Prism 8 (Graph-Pad Software) as indicated in the figure legends. Data are presented as mean ± SEM or ratios among experimental groups and controls ± SEM. When two experimental conditions were compared, statistical significance was calculated by two-tailed t tests. * p < 0.05, ** p < 0.005, ***p < 0.0001.

